# Evolutionary analysis of THAP9 transposase: conserved regions, novel motifs

**DOI:** 10.1101/2021.08.01.454642

**Authors:** Richa Rashmi, Chandan Nandi, Sharmistha Majumdar

## Abstract

THAP9 is a transposable element-derived gene that encodes the THAP9 protein, which is homologous to the *Drosophila* P-element transposase (DmTNP) and can cut and paste DNA. However, the exact functional role of THAP9 is unknown. Here, we perform evolutionary analysis and extensive *in silico* characterization of THAP9, including predicting domains and putative post-translational modification sites. We predict previously unreported mammalian-specific post-translational modification sites that may play a role in the subcellular localization of THAP9. We also observe that although THAP9 has evolved under a strong pervasive purifying selection, yielding high conservation of THAP9, there are distinct class-specific conservation patterns of key functional residues in certain domains. Furthermore, investigation of THAP9 expression profiles in various cancer and matched normal datasets demonstrated underexpression and overexpression in testicular cancers and thymic epithelial tumors, respectively, thus suggesting a possible role of THAP9 in cancer.

## Introduction

Transposable elements (TEs) are DNA sequences that can move and duplicate within a genome^1^. They can cause various mutations as well as an increase in genome size^2^. Thus, TEs, which constitute 25% - 50% of various genomes, play a significant role in evolution.

Many genes are derived from transposable elements. Human THAP9, which encodes the hTHAP9 protein, is one such gene. The hTHAP9 protein is a homolog of the Drosophila P-element transposase (DmTNP) (more than 25% homology)^3^. The DmTNP protein mobilizes the P-element transposon, which is the causative agent for hybrid dysgenesis in Drosophila^4^.

hTHAP9 belongs to the THAP (Thanatos-associated protein) protein family in humans, with twelve proteins (hTHAP0-hTHAP11). All human THAP proteins are characterized by an amino-terminal DNA-binding domain called the THAP domain, which is typically 80-90 amino acid residue-long and possesses a C2CH type Zinc Finger^4, 5^.

Many THAP family proteins are known to be involved in various human diseases. THAP1 is involved in torsional dystonia and hemophilia^6^, THAP5 is involved in heart diseases^7^, THAP2^8^, THAP10^9^, and THAP11^10^ are involved in multiple cancers. Also, the THAP9-AS1 gene, which is the neighboring gene in the 5’ upstream region of THAP9, is overexpressed in pancreatic cancer^11^. Moreover, according to NCBI & the Protein Atlas, THAP9 has high expression in the brain, ovary, prostate, testis & thyroid (https://www.ncbi.nlm.nih.gov/gene/79725, https://www.proteinatlas.org/ENSG00000168152-THAP9/tissue). Nevertheless, the exact functional role of THAP9 and its association with various diseases remains unknown.

Phylogenetic analysis and evolutionary tests of hTHAP9 can help us understand the evolution and the type and extent of natural selection for the hTHAP9 gene and its associated protein. In the course of evolution, negative or purifying selection is used to eliminate unfavorable and harmful mutations^12^. On the other hand, positive or diversifying selection leads to molecular adaptation by promoting favorable mutations^12^. Here, we present the evolutionary analysis and extensive *in silico* characterization, including predicting domains and putative post-translational modification sites for THAP9 and its orthologs. This study identified some previously unreported functional features in the THAP9 protein sequence, highly conserved in mammals. These include four adjacent motifs: N-glycosylation site, Protein kinase C (PKC) phosphorylation site, Leucine zipper domain, and Bipartite nuclear localization signal (NLS), which may play a role in the subcellular localization of THAP9. The study also revealed two N-myristoylation sites within the THAP domain. Moreover, it was observed that THAP9 has evolved under a strong pervasive purifying selection.

## Materials and Methods

### Identification of orthologs

THAP domain containing 9 (*THAP9*) transcript variant 1 (RefSeq: NM_024672.6) from *Homo sapiens* was used to identify the orthologs from the Eukaryotic Genome Annotation pipeline of the NCBI database (National Center for Biotechnology Information, https://www.ncbi.nlm.nih.gov/) using a combination of protein sequence similarity and local synteny information for the available sequences. *hTHAP9* orthologs are present in 216 organisms, including 74 birds, 4 alligators, 7 turtles, 6 lizards, 4 amphibians, 120 mammals, and 1 lamprey. We used the following five representative classes: amphibians, hyperoartia, reptiles, birds, and mammals for comparison and representation purposes.

### Sequence alignment & Phylogenetic analysis

A phylogeny-aware sequence alignment was performed for amino acid sequences using the CLUSTALW^13^ method present in MEGAX^14^ software using the guide trees generated from TimeTree^15^ (http://www.timetree.org/). We created multiple sequence alignments (MSA) separately for all vertebrate classes (birds, alligators, turtles, lizards, amphibians, and mammals) and combined all species. Following the protein MSA of the orthologs, the Coding DNA Sequences (CDS) were codon aligned using PAL2NAL^16^ (http://www.bork.embl.de/pal2nal/). Divergent sequences which created alignment errors were identified and removed manually. The codon alignments and the protein alignments were used to generate the maximum-likelihood tree using MEGAX with 100 bootstrap replicates, and the tree was visualized using iTOL^17^.

### Analysis for Selective Pressure

To characterize the selection patterns on the alignments of THAP9, we used various tools from the HyPhy package^18^ implemented on the Galaxy webserver^19^. With the user-friendly graphical interfaces, new fast methods, and multi-tasking abilities, HyPhy provides better analysis power to users than the PAML^20^ software package. The Galaxy web-server version of HyPhy provides additional advantages (e.g., it does not require any local installation, and we can automate the whole process by creating a workflow). The approximately-maximum-likelihood phylogenetic trees generated using FASTTREE^21^ also implemented on Galaxy webserver were selected with the HyPhy model selection tool and the codon alignment file. For the HyPhy software package analysis, the default set parameters of p-values/posterior probabilities were considered. FEL^22^ (Fixed Effect Likelihood) and FUBAR^23^ (Fast Unbiased Bayesian AppRoximation) were utilized to identify sites evolving under pervasive diversifying and pervasive purifying selection pressures. MEME^24^ (Mixed Effects Model of Evolution) was used to identify episodically diversifying sites. FEL is an ML-based model which assumes constant selection pressure on each site. The FUBAR model estimates site-by-site ω values while MEME allows the distribution of ω to vary over sites and additionally from branch to branch. Significance was assessed by posterior probability > 0.9 (FUBAR) and p-value < 0.1 (MEME) and p-value < 0.1 (FEL)^25^.

### Protein sequence characterization

The domain architecture of hTHAP9 and orthologs was predicted using SMART^26^ (Simple Modular Architecture Research Tool, http://smart.embl-heidelberg.de/) in “batch mode” considering the option for including Pfam^27^ domains. ELM^28^ Prediction tool (Eukaryotic Linear Motif, http://elm.eu.org/) was used to identify short linear motifs using the fasta file containing hTHAP9 & the ortholog protein sequence as input. Furthermore, the previously unidentified conserved motifs and functional domains were searched using Meme-Suite^29^ and ScanProsite^30^, respectively. Disordered binding regions (DBR)^31^ and secondary structure elements of the hTHAP9 protein were predicted using PSIPRED Workbench^32^, and areas with disorder scores greater than 0.6 were considered as disordered. The physiochemical attributes of the hTHAP9 protein sequence such as molecular weight, theoretical pI, amino acid composition, atomic composition, extinction coefficient, estimated half-life, instability index, aliphatic index, and grand average of hydropathy (GRAVY) were computed using Expasy ProtParam^33^ and EMBOSS Pepstats^34^ (https://www.ebi.ac.uk/Tools/seqstats/emboss_pepstats/). We also analyzed the evolution of some previously reported functional domains and residues of hTHAP9 protein sequence individually, which includes THAP-domain, Leucine Zipper domain, Insertion domain & putative Catalytic Residues. Finally, previously annotated SMART & Pfam domains and newly predicted domains of hTHAP9 and its orthologs were visualized using iTOL^17^.

### Distribution of SNPs and mutations in cancer

The 1000 Genomes project^35^ data using Ensembl^36^ (reference genome: GRCh38.p13) was searched for missense mutations of hTHAP9 (ENSG00000168152). PolyPhen-2^37^ scores given on the Ensembl result table were used to classify the SNPs into “benign,” “possibly damaging,”, “probably damaging”. PolyPhen-2 predicts the effect of an amino acid substitution on the structure and function of a protein using sequence homology, Pfam annotations, 3D structures from PDB were available, and several other databases and tools (including DSSP, ncoils, etc.)^38^. Cancer-related missense mutations in THAP9 were identified using cBioPortal^39^ for Cancer Genomics (http://www.cbioportal.org/). PolyPhen2 scores for these mutations were calculated by submitting their genomic coordinates (for GRCh37 reference genome) to the Ensembl Variant Effect Predictor tool^40^ (https://www.ensembl.org/Tools/VEP).

### Cancer expression databases and web tools

To get an idea about the expression profile of the *hTHAP9 gene* in various cancers, we used the expression data from TCGA^41^ (https://cancergenome.nih.gov) and GTEx^42^ (https://gtexportal.org/home). Cancer and matched normal datasets from the above databases were accessed and plotted using the GEPIA^43^ web server (http://gepia.cancer-pku.cn).

## Results

### Evolution of THAP9 through organisms

To study the possible functional role of hTHAP9, we conducted an extensive characterization of its protein sequence and investigated its evolution. hTHAP9 protein is encoded by the gene with the same name and is found on chromosome 4 in humans. Transcript variant 1 of hTHAP9 is known to encode the transposase protein^44^. Therefore, we used the same transcript variant for our analysis.

According to NCBI (Jan 2021), hTHAP9 protein has orthologs in 216 vertebrate organisms, including 74 birds, 4 alligators, 7 turtles, 6 lizards, 4 amphibians, 120 mammals, and 1 lamprey. We downloaded the protein & coding DNA sequences for the same from NCBI. We began our analysis by aligning the protein sequences using CLUSTALW^13^ in a phylogeny-aware manner. We created the time tree for the organisms using timetree^15^. Post-alignment, we filtered the diverse, low-quality, and partial sequences, after which we were left with 178 sequences (3 amphibians, 1 hyperoartia, 13 reptiles, 53 birds,108 mammals). Later we realigned the sequences using CLUSTALW^13^, followed by building Maximum likelihood Phylogenetic trees^45^ with 100 bootstrap replicates. Both these parts were performed using options available in MEGAX^14^. We codon aligned the coding sequences from the orthologs using PAL2NAL^16^, taking corresponding aligned protein sequences as input. The resulting trees (**Figure 1**) were drawn using the iTOL^17^ web server. Rooting trees to the midpoint perfectly aligns the lamprey (hyperoartia), amphibians, reptiles, birds, and mammals in the given order, thus suggesting that THAP9 evolved with the evolution of the organisms, both at the level of DNA (**Figure 1b**) as well as the corresponding amino acid sequence (**Figure 1a**) of the gene.

**Figure 1:**
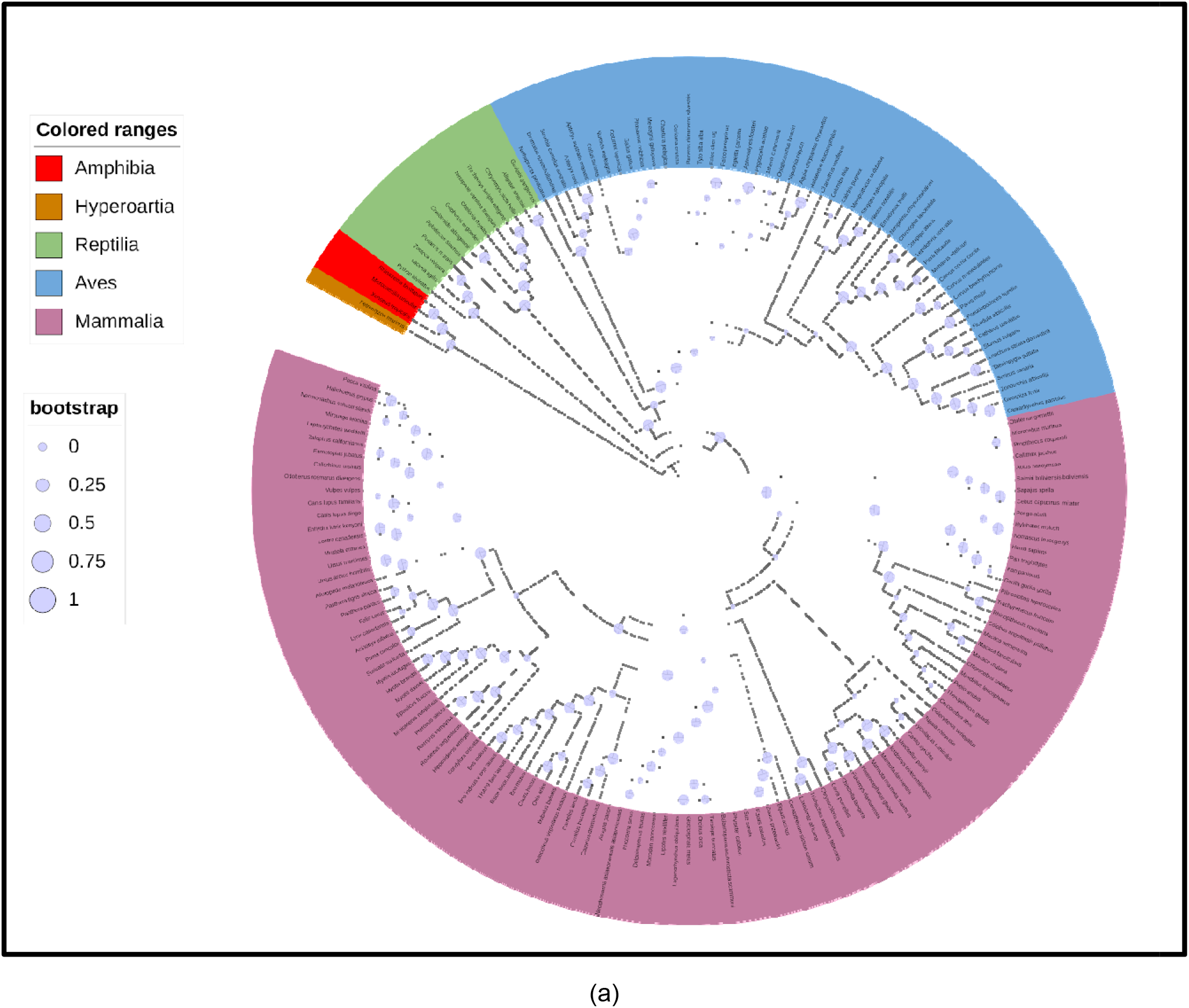
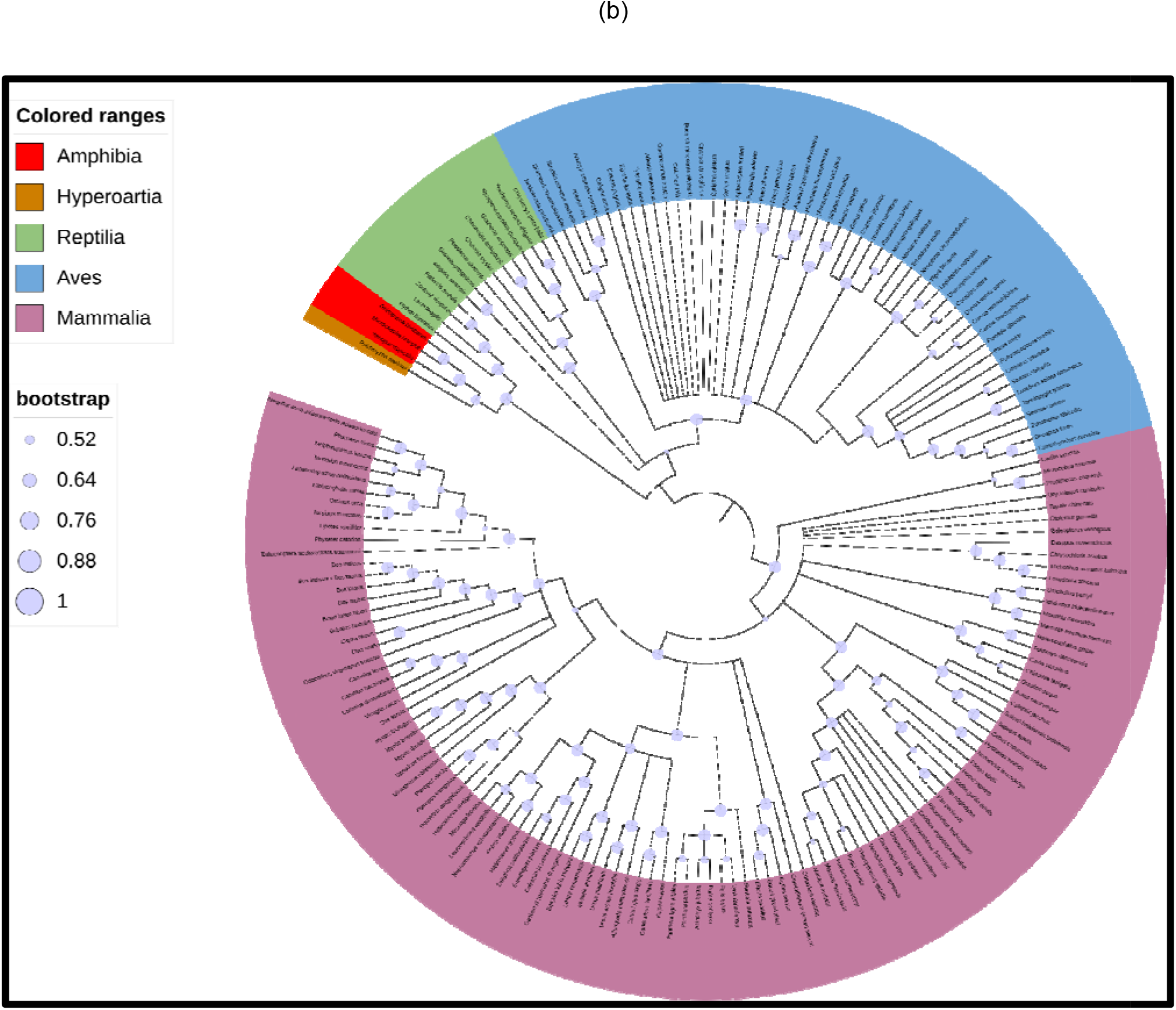
Phylogenetic analysis of THAP9 orthologs. **(a) Phylogenetic tree of aligned protein sequences of THAP9 orthologs.** Protein sequences were aligned using CLUSTALW^13^, followed by building Maximum likelihood Phylogenetic trees^45^ with 100 bootstrap replicates. **(b) Phylogenetic tree of aligned coding DNA sequences of THAP9 orthologs.** Coding DNA sequences of the THAP9 orthologs were codon aligned using PAL2NAL^16^ and Maximum likelihood. Phylogenetic trees were generated using MEGAX^14^ with 100 bootstrap replicates. The trees were generated using iTOL^17^, and the organism classes are marked in different colors (legend on the left). Bootstrap values are marked on the tree.

Several types of tests can be used to speculate about the type and extent of natural selection. Negative or Purifying selection is used to eliminate unfavorable or deleterious mutations. It is mostly observed in housekeeping and essential genes. On the other hand, positive or diversifying selection favors mutations required for molecular adaptation and is mostly observed in genes involved in immunity, reproduction, and sensory systems. Random mutations or selective pressure that mutates gene sequences can cause either silent/ synonymous or non-synonymous mutations^46^ due to degeneracy of the genetic code^51^. The rates of these two types of mutations are denoted as dS and dN, respectively, and their ratio is denoted as (omega); =dN/dS. Thus the value acts as an indicator of how natural selection has acted on the protein. Sites under diversifying/positive selection have a higher rate of non-synonymous mutations and have a value greater than 1. If the site has a higher rate of synonymous mutations, it is under purifying/negative selection and bears a value less than 1. The tools used in our analysis used mathematical models to measure the values for each codon site across the length of the coding DNA sequence by comparing it with the orthologs^47^.

To explore variations in selective pressures across THAP9, we used the HyPhy software^18^ package available at Galaxy^19^ web-server. Our analysis was aimed at the detection of selection at the individual codon level. We used HyPhy-FEL^22^(Fixed Effect Likelihood), HyPhy-FUBAR^23^ (Fast Unconstrained Bayesian AppRoximation), and HyPhy-MEME^24^ (Mixed Effect Likelihood Model) to the codon-aligned sequence file from PAL2NAL^16^. All three methods employ a distinct algorithmic approach to infer selection. FEL uses maximum likelihood to fit a codon model to each site, thereby estimating a value for dN and dS at each site. FUBAR uses the Bayesian approach to selection inference. It uses *a* priori specified grid of dN and dS values (typically 20 × 20), spanning the range of negative, neutral, and positive selection regimes, whose likelihoods can be pre-computed and used throughout the analysis. MEME employs a mixed-effects maximum likelihood approach, where it infers two ω rate classes for each site and corresponding weights representing the probability that the site evolves under each rate class at a given branch^25^.

FEL and FUBAR identified codon sites that evolved under the influence of pervasive (consistently across the entire phylogeny) diversifying and purifying selection, with p-value < 0.1 and posterior probability > 0.9, respectively. FEL predicted 8 diversifying sites and 514 purifying selected sites (**Figure 2 (a)**). Similarly, FUBAR predicted 1 diversifying selected sites and 618 purifying sites (**Figure 2 (b)**). By combining the results, we can say that THAP9 had been subjected to a strong negative selection, with one common positively selected site at Ile829 reported by both FEL and FUBAR.

**Figure 2:**
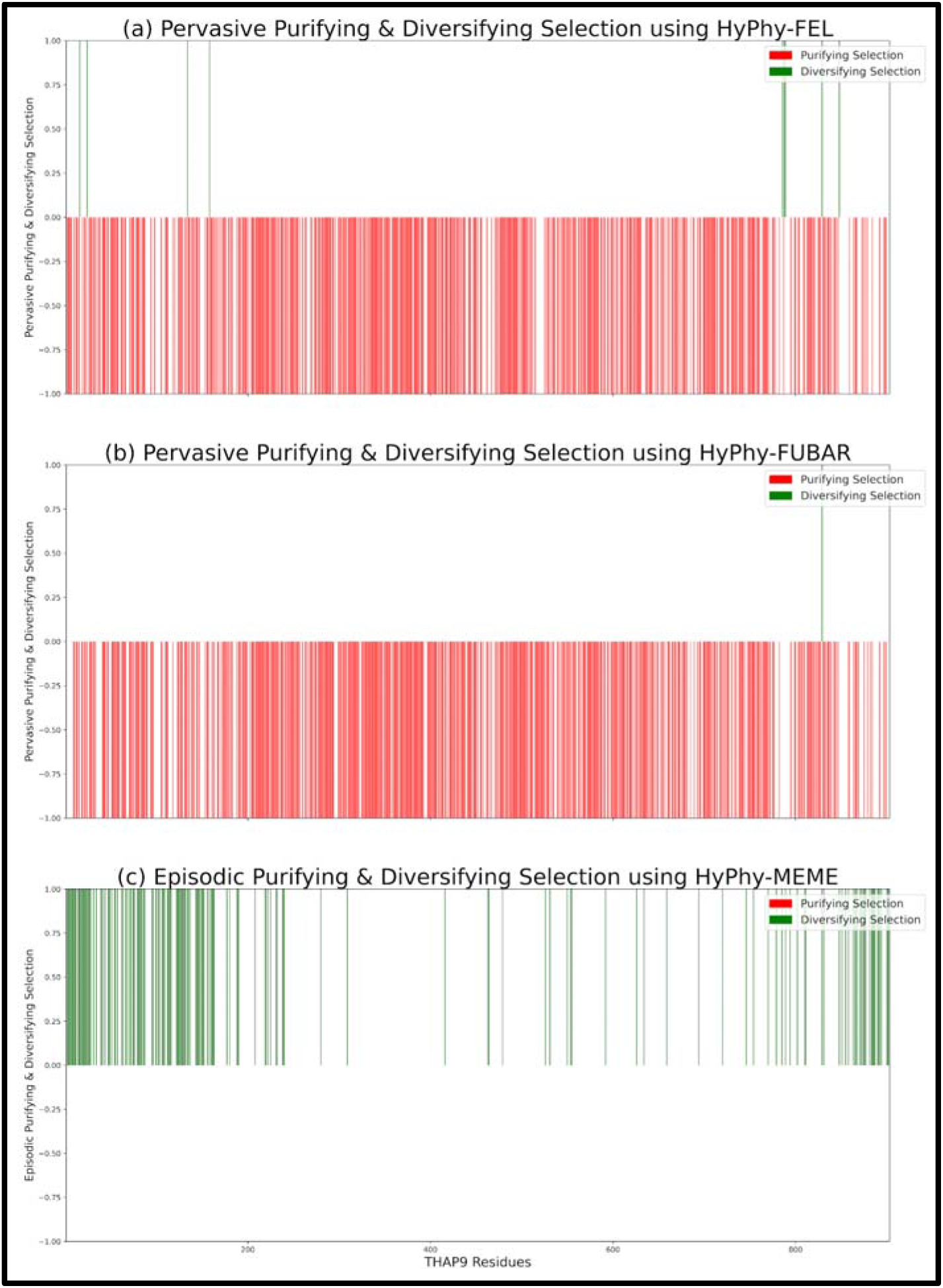
Analysis of Selective Pressure Applied on THAP9 gene. Three independent analyses were performed using HyPhy-FEL^22^, HyPhy-FUBAR^23^, and HyPhy-MEME^24^ packages. Significance was assessed by posterior probability > 0.9 (FUBAR) and p-value < 0.1 (MEME) and p-value < 0.1 (FEL).**(a)** FEL predicted 8 diversifying sites and 514 purifying selected sites. **(b)** FUBAR predicted 1 diversifying selected site and 618 purifying sites. **(c)** MEME observed 197 codon sites, distributed across the hTHAP9, to be evolved under episodic diversifying selection.

MEME identifies site(s) in a gene subjected to both pervasive or episodic (only on a single lineage or subset of lineages) diversifying selection^25^. 197 codon sites (**Supplementary Table 1** https://docs.google.com/spreadsheets/d/1PJQu_fhi2EGv0MyaZHebu1_swTksFNDZG_aEVMTOwgU/edit#gid=1057877600) distributed across hTHAP9 were observed to have evolved under episodic diversifying selection through this analysis (**Figure 2 (c)**).

Based on the results from all 3 methods, we can say that THAP9 has evolved under a strong negative selection suggesting that THAP9 may have some stringent functional or structural requirements. The presence of many episodic positively selected sites suggests that certain essential mutations were favored across some lineages. Moreover, there was only one common positively selected site at Ile829 detected by all three methods.

### Analysis of THAP9 protein orthologs suggests new functional motifs in mammals

The physicochemical properties and domain architecture of THAP9 protein orthologs were predicted using various tools.

#### Physicochemical properties of hTHAP9

We predicted the physicochemical properties using Expasy ProtParam^33^ and EMBOSS Pepstats^34^. hTHAP9 protein is a 903 amino acid long protein with a molecular weight of 103410.79 Da. The isoelectric point (pI) of hTHAP9 is 9.4853. The instability index (Proteins having instability index<40 are considered stable) of hTHAP9 is 38.49. The GRAVY value of hTHAP9 is -0.179, which predicts better interaction between the protein and water. The amino acid composition of hTHAP9 is shown in **Figure 3**.

**Figure 3:**
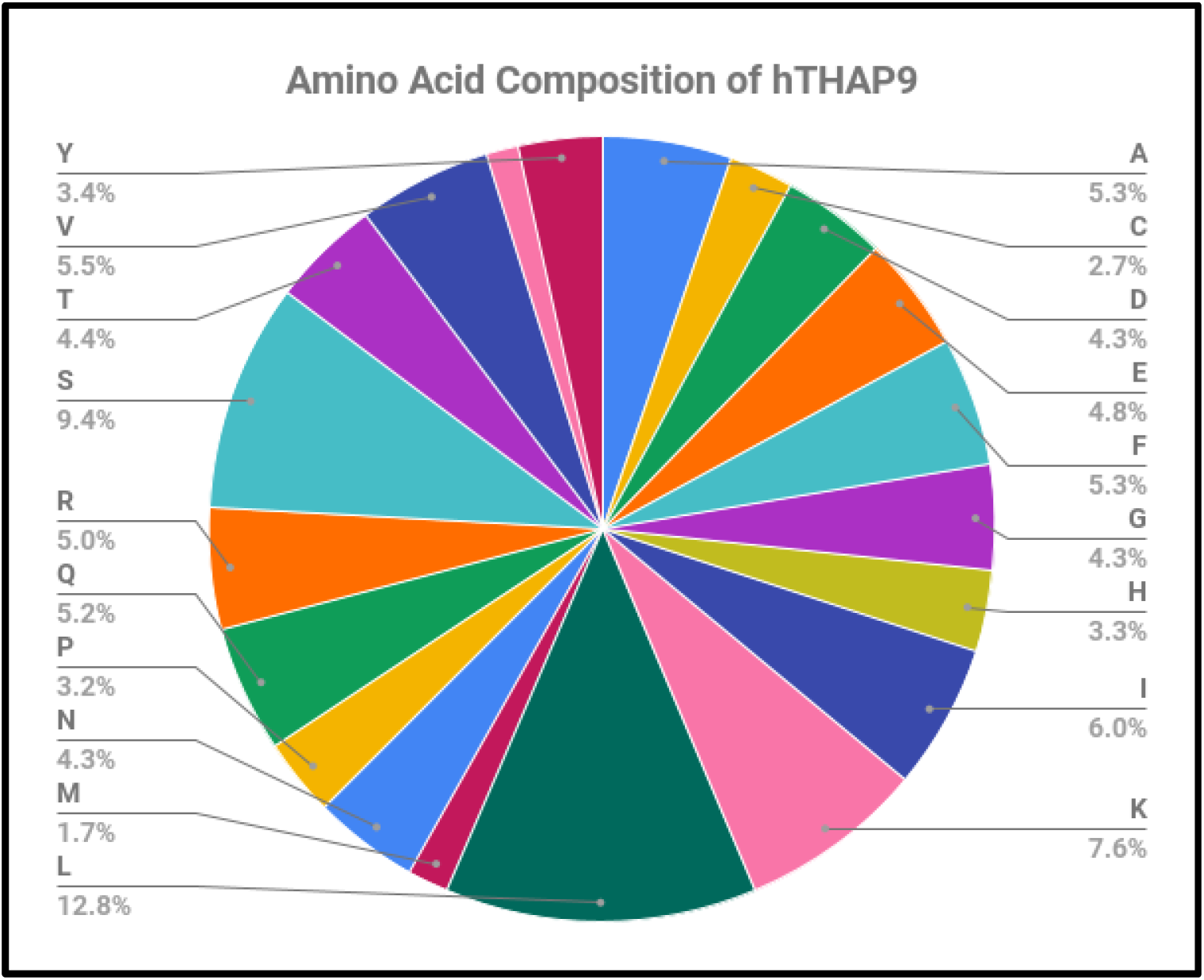
Amino Acid composition of human THAP9 protein.

#### Domain architecture

According to SMART^26^ & Pfam^27^, *hTHAP9* has an 89 residue-long THAP-type Zinc Finger domain at the N-terminal end followed by a 177 residue-long P-element transposase domain (Tnp_P_element, Pfam ID: PF12017) between residues 142 to 389 (**Figure 4**). SMART also predicted two low complexity regions in the h*THAP9* protein sequence from residues 444 to 455 and 567 to 579. Most of the THAP9 orthologs have similar SMART & PFAM annotations. All the annotated THAP9 ortholog sequences have the Tnp_P_element domain. However, some species lack the THAP domain, for example, *Falco Peregrinus* (Aves) and *Alligator Sinensis* (Reptile) (**Figure 4**, full representation in **Supplementary Figure 1(a)**). There are no THAP9 orthologs of Amphibians and Hyperoartia class with already annotated SMART or Pfam domains.

**Figure 4:**
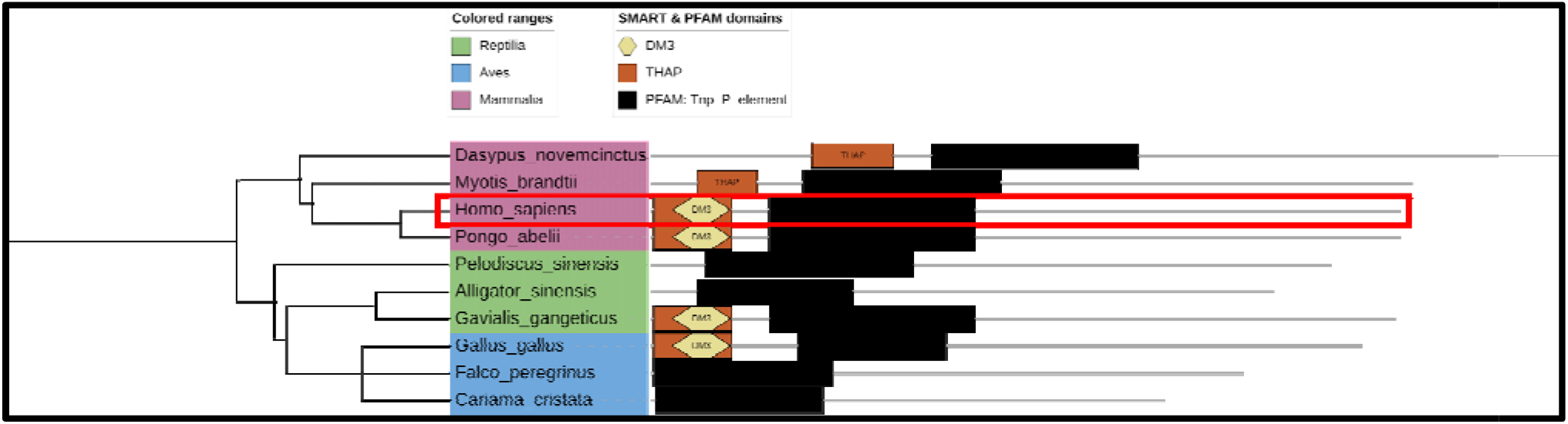
Previously annotated SMART^26^ & Pfam^27^ domains of THAP9 orthologs. Results were visualized on a phylogenetic tree using iTOL^17^. Domain architectures were displayed using the data export feature of SMART.

#### Functional Motifs

Various new functional features of THAP9 were predicted using the Eukaryotic Linear Motif^28^ (ELM) and ScanProsite^30^ (**Figure 5**, full representation in **Supplementary Figure 1(b)**).

**Figure 5:**
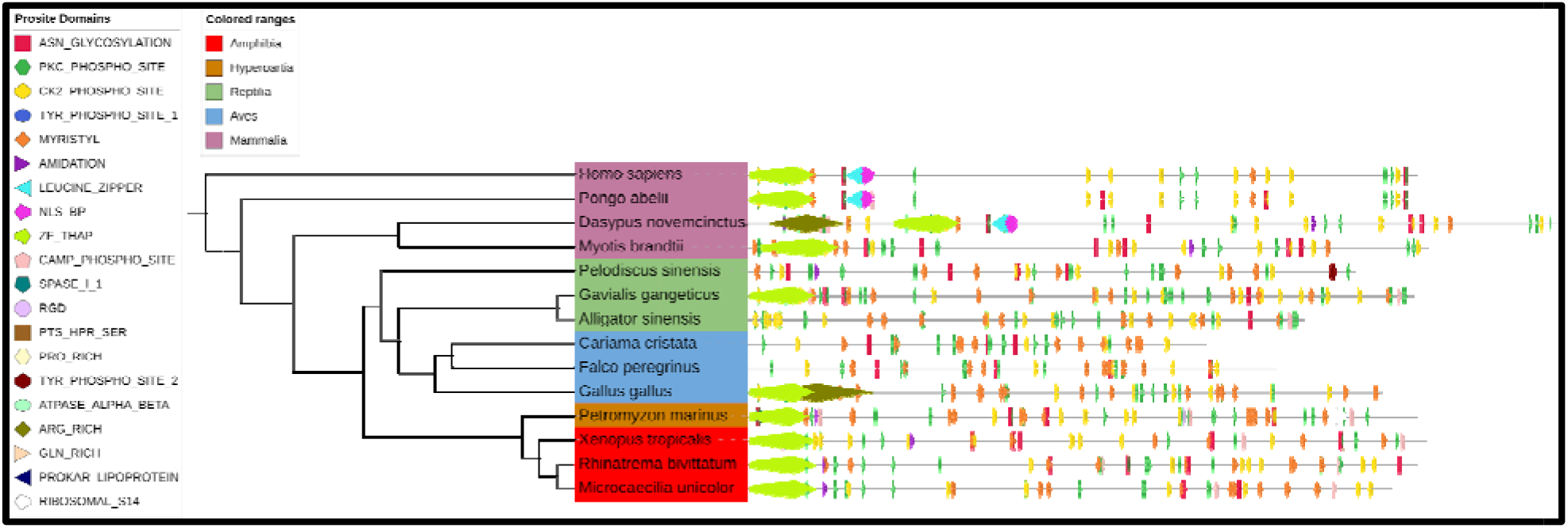
New predicted functional features of THAP9 orthologs. using Eukaryotic Linear Motif^28^ (ELM) and ScanProsite^30^ . The tree was made using iTOL^17^.

3 N-myristoylation sites (Prosite ID: PS00008) were predicted in hTHAP9, namely (i) “GAilCS” located between residues 58 to 63, (ii) “GAvpSV” located between residues 83 to 88, and (iii) “GVsvTK” located between residues 679 to 684. The first two N-myristoylation sites are located in the N terminal THAP domain. hTHAP9 may localize to the nucleus as it has a sequence-specific DNA binding THAP domain^4, 5^ as well as a predicted bipartite NLS^48^ (Figure 6). However, myristoylation is a protein-lipid modification essential for cellular signaling, protein-protein interactions as well as intracellular targeting of proteins to endomembrane or plasma membrane systems^49^. Thus, it is possible that THAP9 could also localize to cytosolic locations via selective myristoylation.

**Figure 6:**
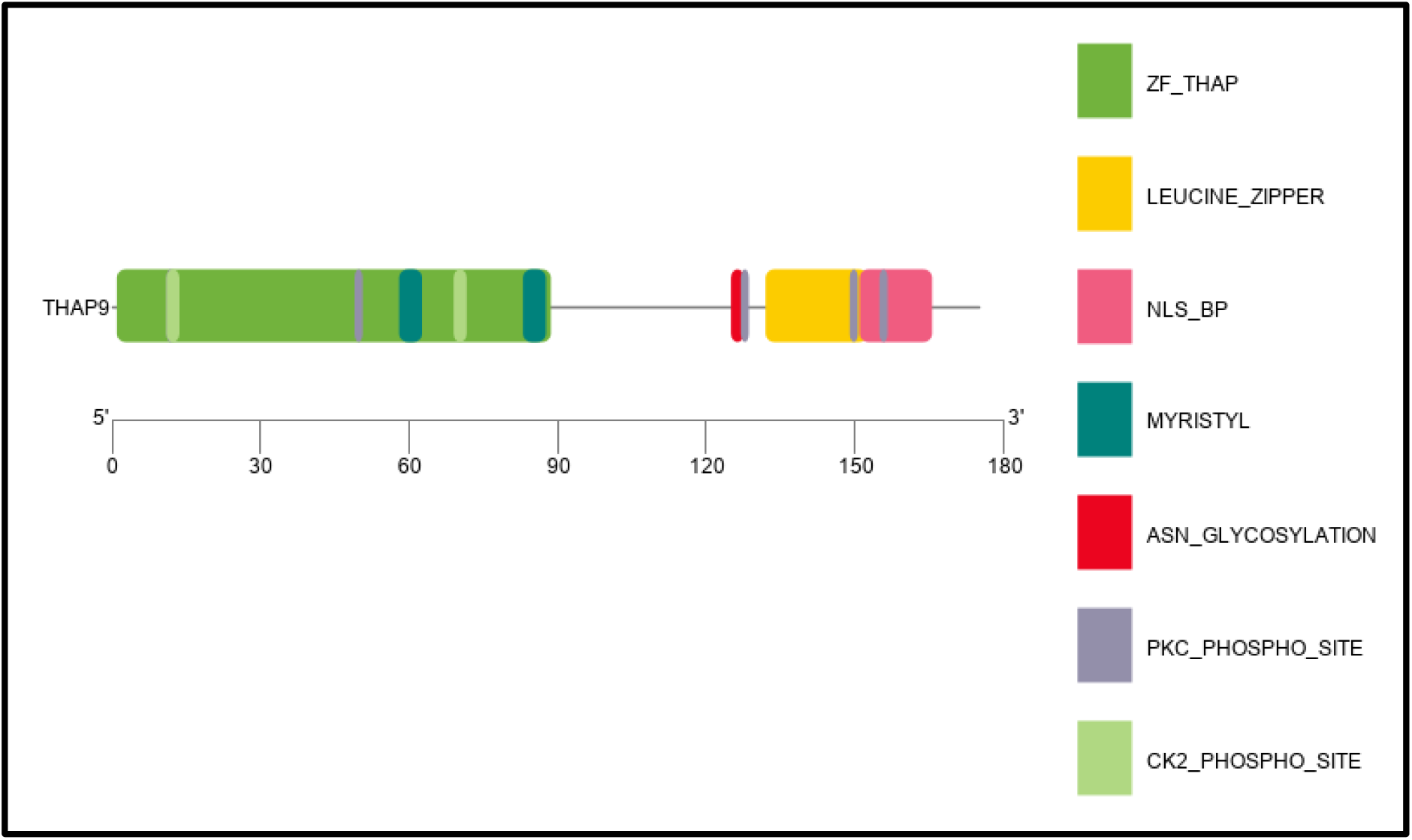
Architecture of N terminal (1-175) region hTHAP9. Predicted N-myristoylation sites (PS00008) (i) “GAilCS” (residues 58-63), (ii) “GAvpSV” (residues 83-88), and (iii) “GVsvTK”(residues 679-684). Predicted 4 adjacent motifs (i) “NYSL” N-glycosylation site (PS00001) (125-128) (ii) “SlK” Protein kinase C phosphorylation site (PS00005) (127-129) (iii) “LtigaekLaevqqmLqvskkrL” Leucine zipper pattern (PS00029) (132-153) (iv) “KRLISVKNYR-MIKKRK” Bipartite nuclear localization signal profile (PS50079) (151-166). This figure has been created using TBtools^51^.

Moreover, the ELM and ScanProsite analysis also predicted the occurrence of 4 adjacent motifs located between 125 to 166, namely (i) “NYSL” N-glycosylation site (PS00001) from 125 to 128 (ii) “SlK” Protein kinase C phosphorylation site (PS00005) from 127 to 129 (iii) “LtigaekLaevqqmLqvskkrL” Leucine zipper pattern (PS00029) from 132 to 153 (iv) “KRLISVKNYR-MIKKRK” Bipartite nuclear localization signal profile (PS50079) from 151 to 166. This pattern is highly conserved in mammals (**Figure 6, Supplementary Figure 1(b)**).

Multiple previously unreported functional motifs (**Supplementary Table 2**) were also predicted. These motifs can be explored further to understand the regulation and function of THAP9.

#### NLS

NLS sequences generally appear either as a single-stretch (monopartite) or as two clusters (bipartite) of basic residues separated by approximately ten amino acid residues, with the respective consensus sequences being (K/R)4–6 and (K/R)2 X10–12(K/R)3^50^. ELM and ScanProsite analysis found a 16 residue long bipartite NLS sequence “KRLISVKNYR-MIKKRK,” in which 50% of the residues were basic, and the basic region was more concentrated in the C-Terminal end of the motif.

Bipartite nuclear localization signals (NLS) are sometimes located close to (e.g., HSF2^51^) or within (e.g., SREBP-2 ^52^) a Leu-zipper domain. Moreover, the subcellular localization of a protein may be regulated by masking and unmasking of its NLS by local phosphorylation events^52^. It is tempting to speculate that the subcellular localization of hTHAP9 may be regulated by its Leucine zipper domain which contains a bipartite NLS as well as a highly conserved Protein kinase C phosphorylation site.

#### Phosphorylation sites

Several kinase phosphorylation sites (Protein kinase C & Casein kinase II) were predicted to be distributed across the length of the protein. ELM also predicted one Host Cell Factor-1 binding motif in hTHAP9 that has been previously annotated^28, 48^.

#### Other domains

When we looked for weak domain matches in THAP9, ScanProsite also predicted the presence of a Phosphatase tensin-type domain profile (PPASE_TENSIN). Tumor suppressor protein PTEN is the best-characterized member of the PPASE_TENSIN family^53^. To further explore the weak presence of the PPASE_TENSIN (Prosite Entry: PS51181) domain in the human THAP9 protein, we looked for this domain in all the THAP9 ortholog sequences using ScanProsite (Option 3). The PPASE_TENSIN domain was identified in 105 orthologs.

The Phosphatase tensin-type domain is present in 38 protein sequences in the Uniprot database^54^, including human PTEN. To look for conserved motifs in the predicted PPASE_TENSIN domain sequences in 105 THAP9 orthologs and the 38 proteins from Uniprot, we processed them through the MEME tool part of the MEME-Suite^29^ using site distribution parameter as “one occurrence per sequence.” We identified 15 highly conserved motifs, but the confidence score of these matches was not significant enough.

To further examine the diversification of THAP9 orthologs, we predicted conserved motifs using MEME^29^. We identified the distribution of the top 30 conserved motifs occurring at least once in all the sequences (**Figure 7; Table 1**). The motif composition appeared more conserved in mammals than other groups, suggesting functional relevance in mammals as well as functional diversification among other classes.

**Figure 7:**
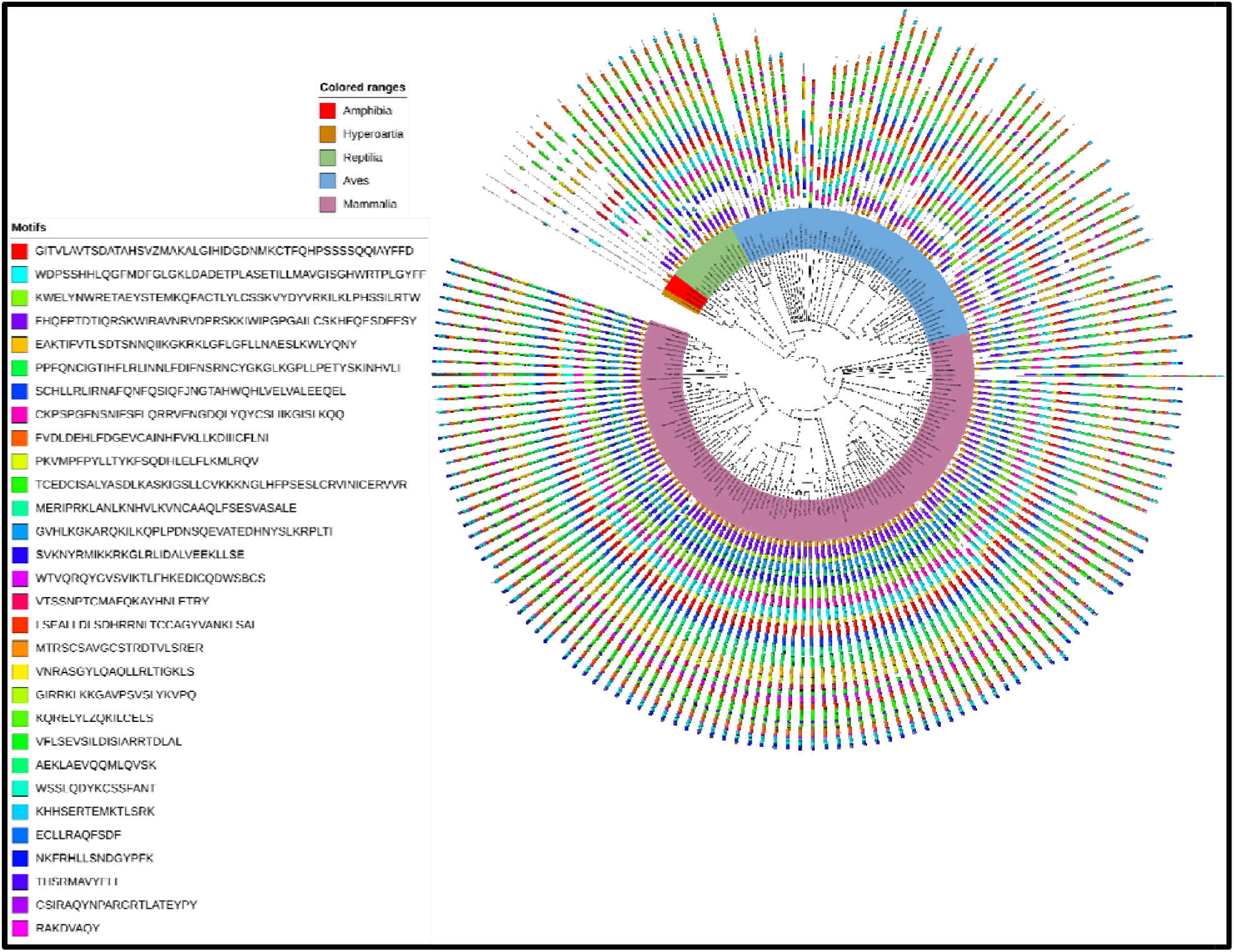
Top 30 MEME Motifs from THAP9 Ortholog Proteins. Schematic distribution of top 30 conserved motifs predicted in protein sequences of THAP9 orthologs. Motif analysis of 173 sequences was performed using MEME-suite^29^. The phylogenetic tree was constructed using the Maximum likelihood method available in MEGAX^45^. The trees were generated using iTOL^17^, and the organisms’ classes are marked in different colors with the legend given on the left side.

### Evolution of individual domains

To study the functional diversification of THAP9 among different vertebrate classes in more detail we studied the evolution of some previously reported functional domains and residues of functional relevance. We looked at the evolution of the THAP domain, L-Zipper domain, predicted insertion domain as well RNaseH-like catalytic domain^55^ (including 8 putative catalytic residues namely K282, D304, D374, D414, D519, E613, D695, E776 in hTHAP9) in 178 orthologs.

#### THAP-Domain

The THAP domain, the characteristic feature of the THAP family proteins, is a conserved 80-90 amino acid DNA binding domain located at the N-terminal end of the protein. The THAP domain is a C2CH type Zinc Finger domain that can bind specific DNA sequences and has a conserved secondary structure, namely a characteristic β-α-β fold with four loops (L1-L4) interconnecting the sheets and the helix^5^.

Our analysis showed that the THAP-Domain is present in all amphibian (3) and hyperoartia (1) orthologs and in most orthologs in reptiles (12 out of 13), birds (26 out of 53), and mammals (105 out 108). The THAP domain shows high conservation within these organism classes (**Figure 8, Supplementary Table 3**). In a few organisms, the THAP domain was truncated (N Terminal deletion: *Apteryx rowi, Piliocolobus tephrosceles, Tupaia Chinensis*; C Terminal deletion: *Haliaeetus leucocephalus, Chinchilla lanigera, Puma concolor*).

**Figure 8:**
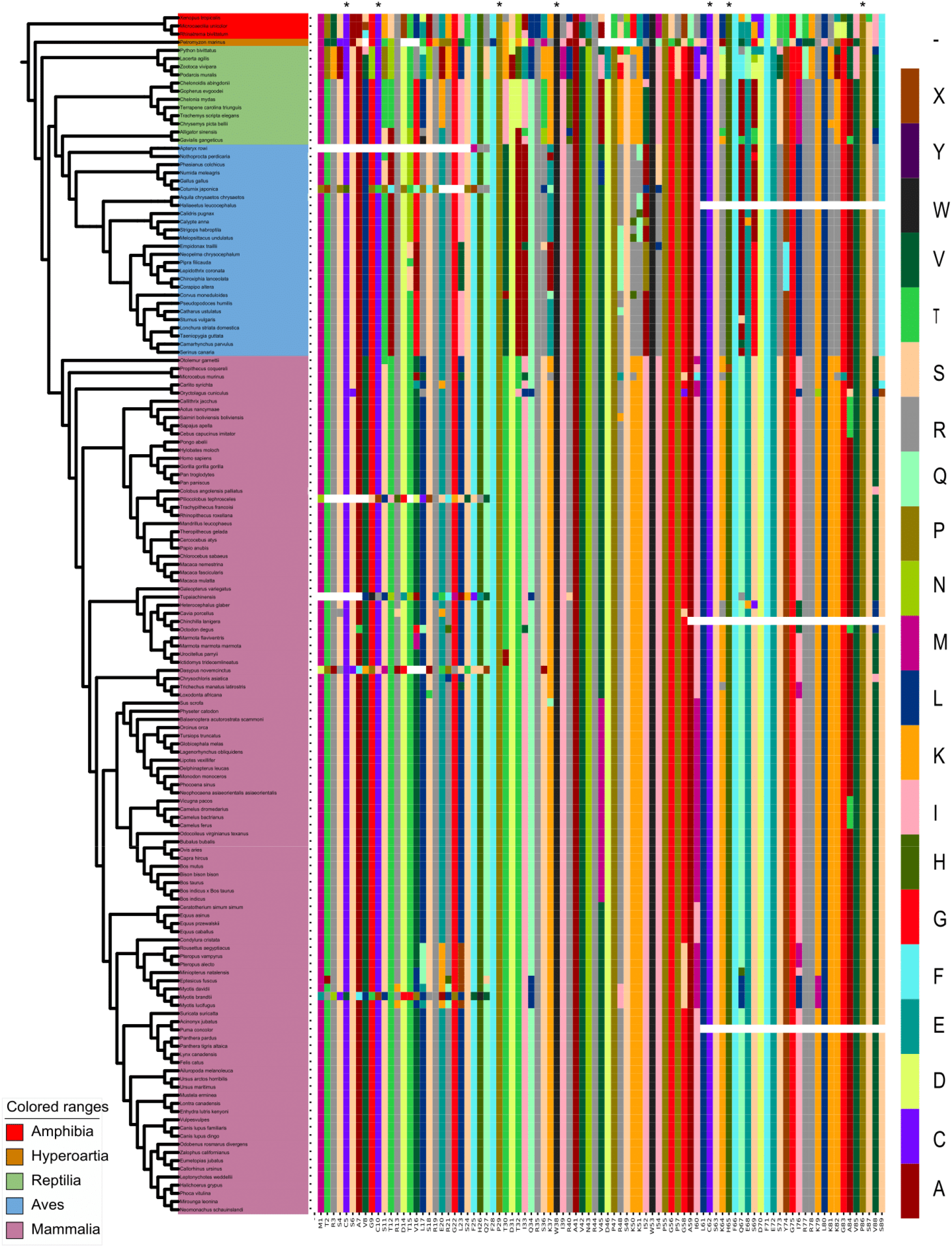
Evolution of THAP domain in THAP9 orthologs. Conserved functional residues (C5, C10, C62, H65; P29, W38, V85, and P86 in hTHAP9) have been highlighted with a * mark at the top.

Structure-based multiple sequence alignment of the THAP domains of DmTNP, human THAP1, 2, 7, 9, and 11, and C. elegans CtBP by Sabogal et al.^4^ reported the conservation of zinc-coordinating C2CH motif and base-specific DNA-binding residues (corresponding hTHAP9 residues are C5, C10, C62, H65, P29, W38, V85, and P86 (* marked in **Figure 8**). We investigated the conservation of these residues across the THAP9 orthologs. It was observed that within the 147 orthologs containing THAP domains, C5 is conserved in 142 orthologs (mutated in 3 mammals and 2 aves), C10 is conserved in 141 orthologs (mutated in 4 mammals and 2 aves), C62 is conserved in 144 orthologs (mutated in 2 mammals and 1 aves), H65 is conserved in 143 orthologs (mutated in 3 mammals and 1 aves), V85 is conserved in 137 orthologs (mutated in 2 mammals, 1 aves, 4 reptiles, and 3 amphibians), P86 is conserved in 144 orthologs (mutated in 2 mammals and 1 aves). Interestingly, P29 and W38 are conserved in all 147 orthologs. Moreover, the THAP domain showed a distinct pattern within bats, camels, whales, rodents & lizards (**Supplementary Table 3**).

#### Leucine Zipper Domain

hTHAP9 has a ∼ 40 amino acid long leucine-rich region located downstream of the DNA-binding THAP domain^56^. This was corroborated by our ScanProsite analysis which also predicted a highly conserved L-Zipper domain in the same region. Several DNA transposases like DmTNP^57^, IS911^58^ and KP repressor^59^ (inhibitor of DmTNP) form multimers using the leucine zipper region. In another study, it was observed that mutating the leucines (L90, L128, L132, L139, L146, and L153 to Ala, either individually or together) or deleting the leucine-rich predicted coiled-coil region in hTHAP9 did not disrupt homo-oligomerization^48^. This suggests that maybe THAP9 utilizes a multidomain mechanism for oligomerization. So, we decided to look at the evolution of the L-Zipper domain of THAP9 to get a further understanding of its conservation.

We observed that the L-Zipper Domain is not present in any of the amphibians, hyperoartia, and reptilian orthologs of THAP9. Moreover, it is only present in 14 out of 53 Aves orthologs and 91 out of 108 mammalian orthologs. Thus, it is interesting to note that the L-Zipper domain appeared much later during vertebrate evolution and is highly conserved across mammals. Moreover, L132, L139, L146, and L153 (residue numbers correspond to hTHAP9, **Supplementary Table 4,** highlighted with * in **Figure 9**) are conserved in all the orthologs in which L-Zipper is detected.

**Figure 9:**
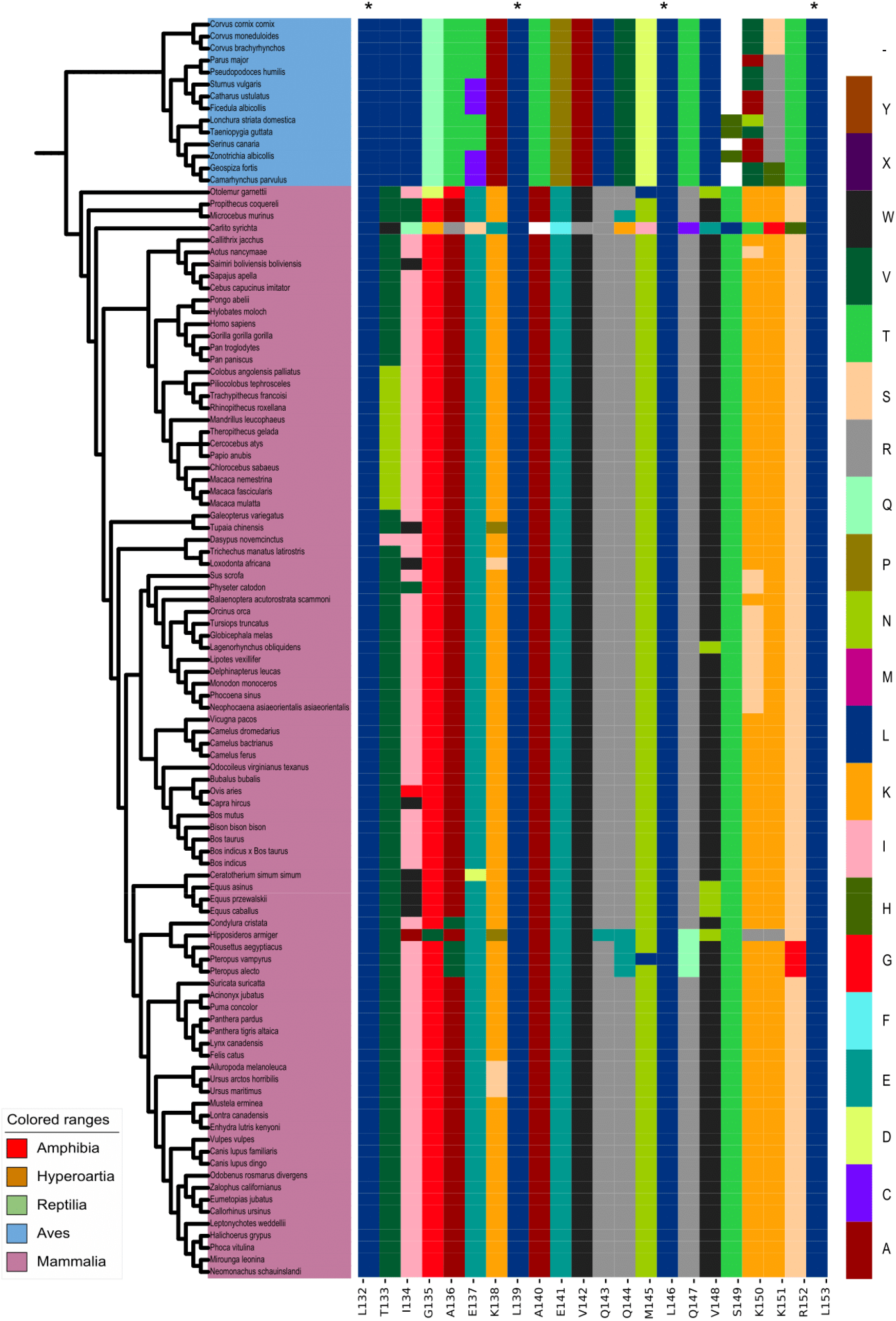
Evolution of Leucine Zipper Domain in THAP9 orthologs. L132, L139, L146, and L153 (residue numbers correspond to hTHAP9) highlighted with *.

#### RNase-H-like fold

A recent study^55^ predicted an RNase-H-like fold in hTHAP9 using secondary structure predictions, homology modeling, and multiple sequence alignment. The RNase-H fold brings three catalytic residues (DDD/E) in close proximity, which coordinate with two divalent metal ion cofactors (mostly Mg2+) to form the active site for DNA cleavage and strand transfer during transposition^60, 61^. RNase-H fold-containing proteins are generally multi-domain in nature with domains involved in sequence-specific DNA binding and multimerization in addition to the RNaseH-like catalytic domain^55, 57, 62^. The study experimentally confirmed that D304, D374, and E613 are important catalytic residues responsible for DNA excision^52^. The catalytic triad of DmTNP is formed by D230, D303 & E531. It was observed that D374 and E613 of hTHAP9 align with D303 and E531 catalytic residues of DmTNP. But D304 of hTHAP9 does not align with D230 of DmTNP, instead, D230 of DmTNP aligns with K282 of hTHAP9. Moreover, they also observed that besides the predicted catalytic triad, there were other conserved acidic residues in the signature string of hTHAP9, which were important for DNA transposition including D414, D519, and D695. They also predicted that the RNase-H fold in hTHAP9 appears to be disrupted by an insertion domain between S415 to T604, similar to the insertion domain of DmTNP^57^ which binds GTP via GTP-binding motifs. Despite the disruption of the RNase-H fold, both DmTNP and hTHAP9 are able to cut the substrate DNA^55, 57^. We investigated the evolution of these important catalytic residues i.e. K282, D304, D374, D414, D519, E613, D695 & E776 (**Figure 10, Supplementary Table 5**). Interestingly K282 was observed to be D in 2 out of 3 Amphibians and D in the hyperoartia. Moreover, out of 13 reptiles, it was A in 1, D in 2, G in 3, and E in 7. But K282 was Q in 51 out 53 birds and K in 107 out of 108 mammals. With this, we can speculate that over the course of evolution the catalytic residue D got changed to K which possibly contributed to the hindered transposition mechanism of hTHAP9 and thus led to the domestication of hTHAP9. Interestingly, while most of the catalytic residues were conserved across all THAP9 orthologs, D374 and D695 were only conserved across the mammalian orthologs and exhibited distinct class-specific substitutions in other classes. It is tempting to speculate that such class-specific conservation patterns of key catalytic residues, play important roles in the modification of THAP9’s function

**Figure 10:**
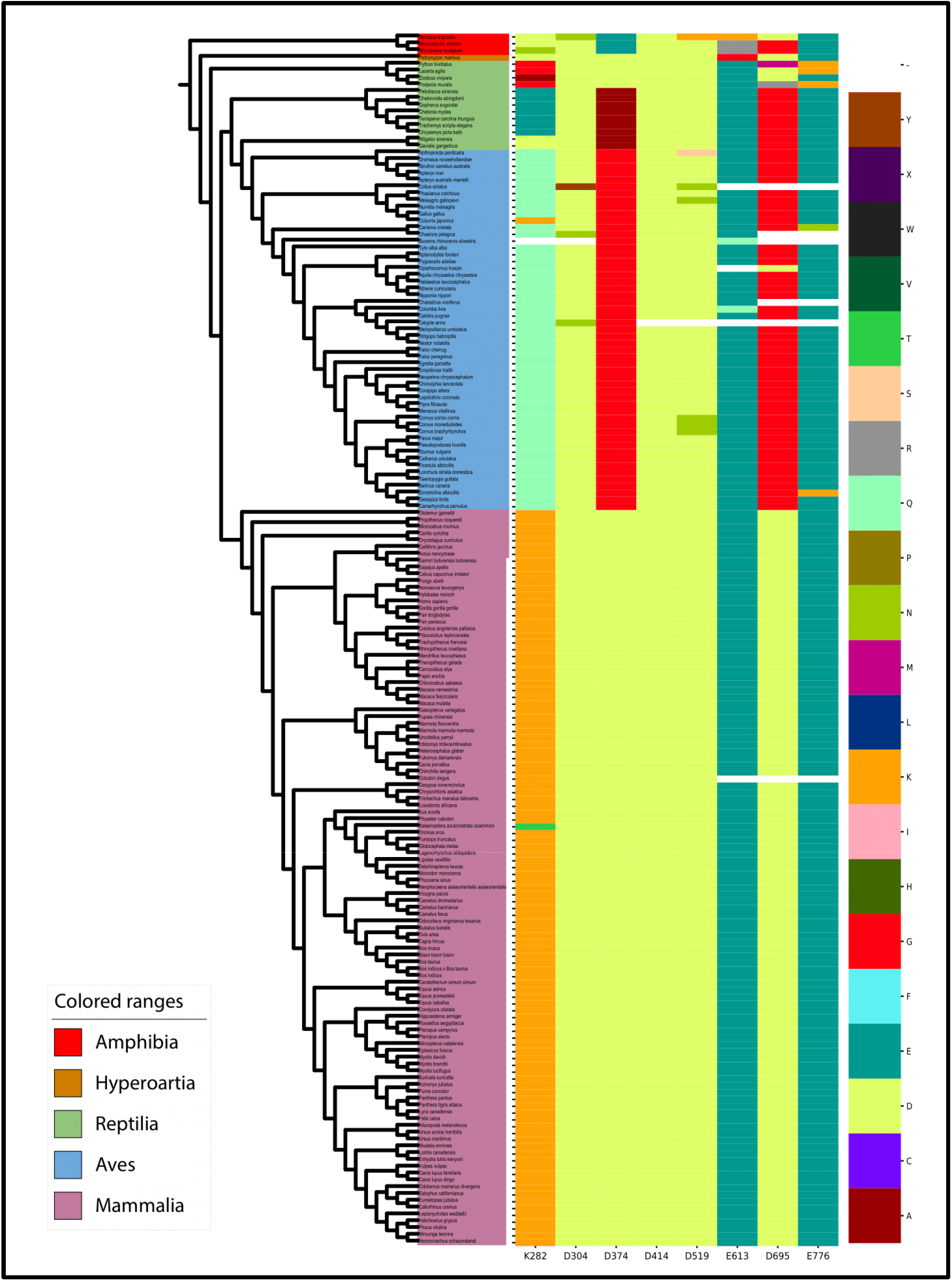
Evolution of catalytic residues of the RNaseH-like catalytic domain in THAP9 orthologs. D304, D374, D414, D519, E613, D695 & E776 are the important catalytic residues of hTHAP9^55^.

The predicted insertion domain, which lies between S415 to T604 of hTHAP9 (**Figure 11, Supplementary Table 6**), exhibited weak conservation across orthologs.

**Figure 11:**
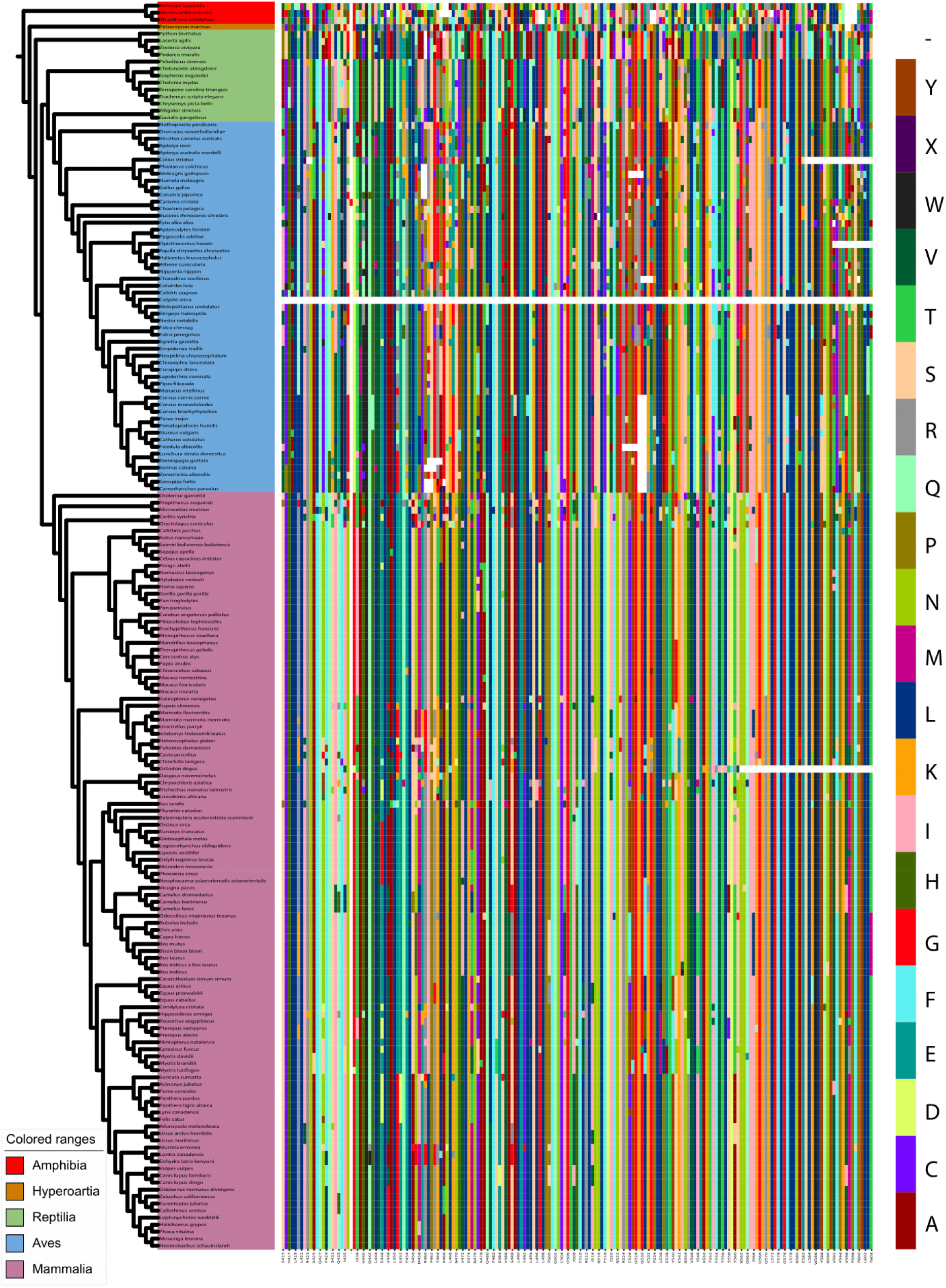
Evolution of predicted insertion domain in THAP9 orthologs.

### Pathogenic mutations in h*THAP9*

We wanted to investigate the possible role of hTHAP9 in disease. No disease-specific mutation has been reported for the hTHAP9 gene (**Supplementary Table 7**). So, we examined the missense SNPs reported in the hTHAP9 gene on the Ensembl Portal^36^ (which uses the 1000 Genomes project^35^ data). PolyPhen-2^38^ scores given on the Ensembl result table classified these SNPs into three categories: 1) score from 0 to 0.444 as “benign,” 2) scores from 0.453 to 0.906 as “possibly damaging,” 3) scores from 0.909 to 1 as “probably damaging.”

According to Ensembl data, ∼79% of missense mutations were “non-pathogenic,” and ∼21% of them were “pathogenic” in nature (**Table 2,** **Figure 12**).

**Figure 12:**
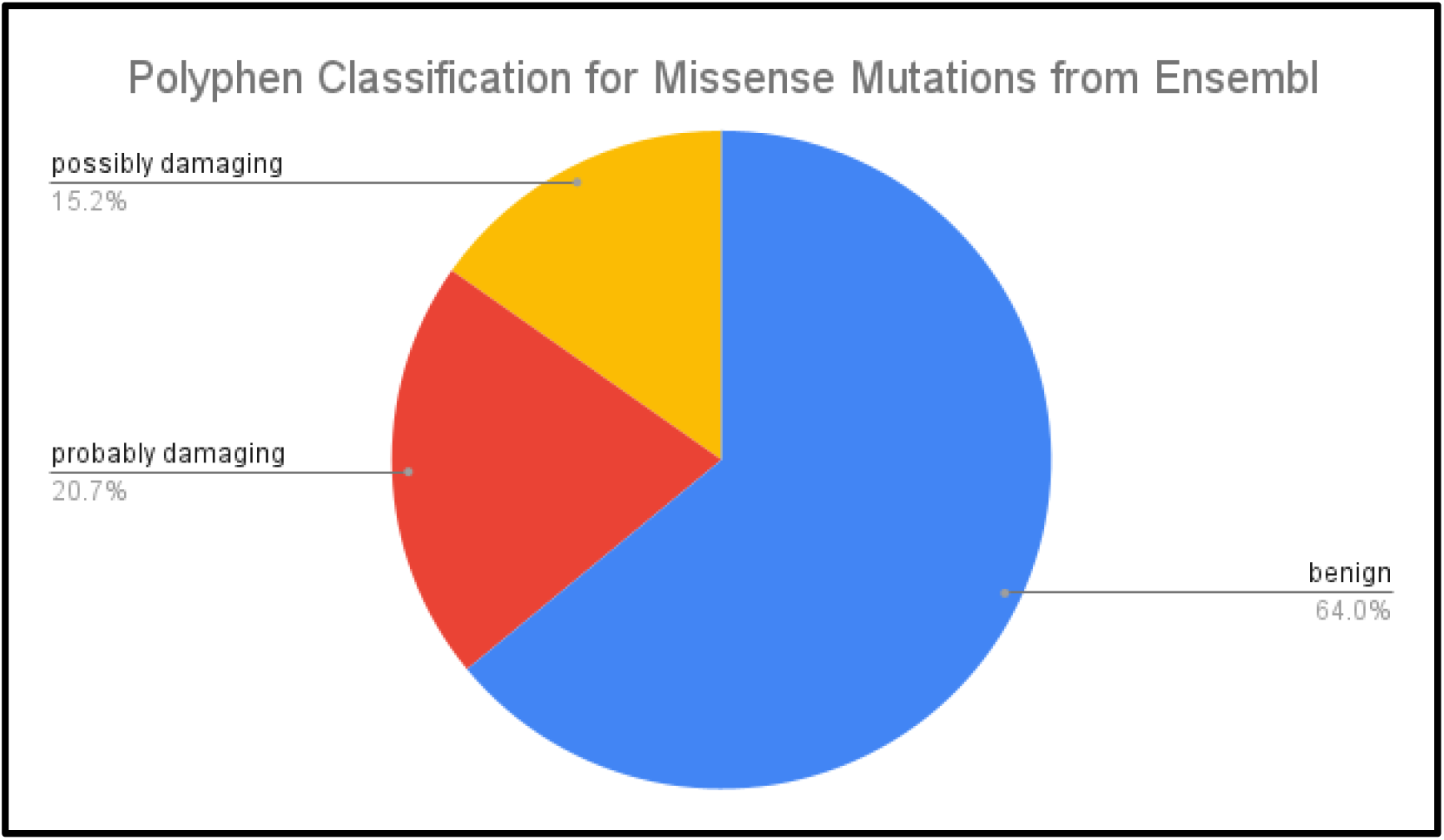
PolyPhen classification for Missense Mutations from Ensembl. PolyPhen classification of 820 Missense SNPs of human THAP9 gene (from Ensembl) into benign, possibly- damaging, and probably damaging categories.

Moreover, we also analyzed the missense mutations found in hTHAP9 in different cancer samples using cBioPortal^39^ (Table 3, **Figure 13**). Following the Ensembl trend,∼74% of missense mutations from CBioPortal were “non-pathogenic,” and ∼27% of them were “pathogenic” in nature. No significant hotspots for mutations in cancer were observed as the mutations were distributed throughout the THAP9 sequence.

**Figure 13:**
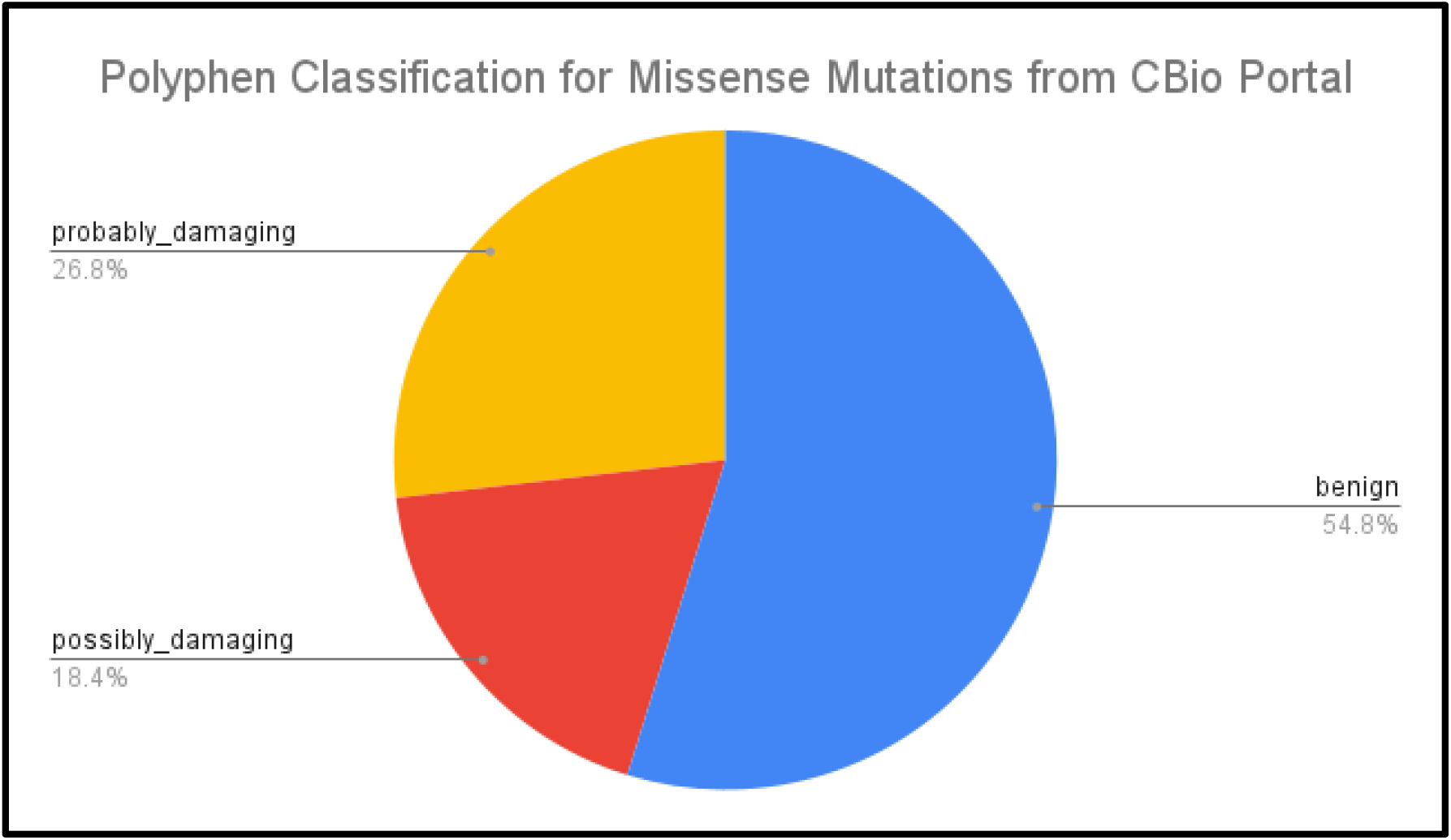
PolyPhen classification for Missense Mutations from CBio Portal. PolyPhen classification of 272 Missense SNPs of human THAP9 gene (from cBio Portal) into benign, possibly- damaging, and probably damaging categories.

The only missense SNP that has a global MAF > 0.05 (minor allele frequency) was rs897945 (MAF = 0.3253), making it a candidate for a derived allele (DA). Derived alleles are new alleles, having a population frequency of at least 5%, that is formed by the mutation of ancestral alleles. It has been reported that the disease-associated alleles are more likely to be low-frequency derived alleles^63^. In the *hTHAP9* gene, rs897945 yields a G → T nucleotide substitution, leading to a Leucine to Phenylalanine mutation at position 299 on Tnp_P_element (Pfam ID: PF12017) domain (**Figure 14**). This SNP may have a role in Atopy and Allergic Rhinitis in a Singapore Chinese Population^64^.

**Figure 14:**
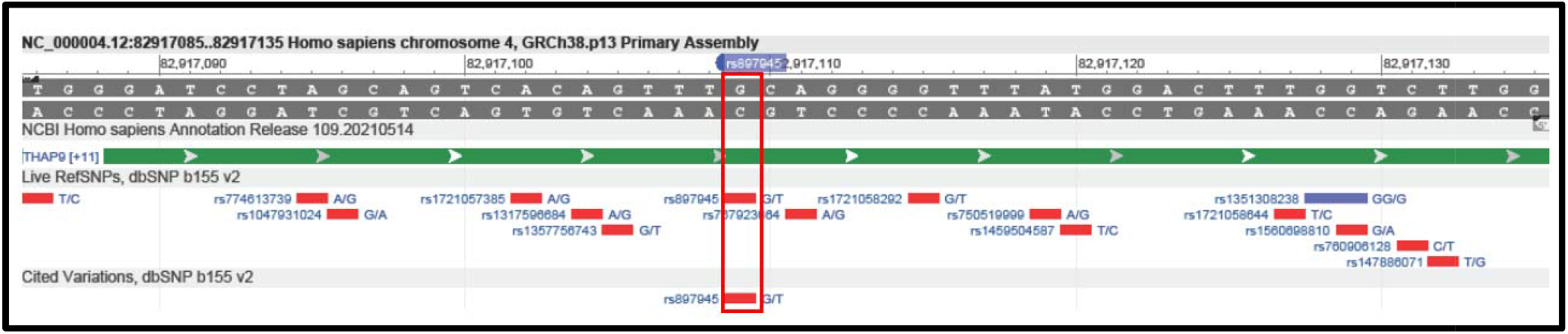
Genome Variation Viewer view of rs897945. rs897945 yields a G → T nucleotide substitution which leads to Leucine to Phenylalanine amino acid change at position 299 located on th Tnp_P_element (Pfam ID: PF12017) domain in hTHAP9 protein.

### *THAP9* shows an altered expression in the case of Testicular & Thymic Epithelial Tumors

To investigate the gene expression profile of hTHAP9 in various cancers, we used the GEPIA^43^ tool, which used cancer and matched normal from TCGA^41^ and GTEx^42^ portal. The results showed overexpression of hTHAP9 in testicular cancer and reduced expression in Thymic Epithelial Tumors (THYM) (**Figure 15**). We can thus speculate that hTHAP9 may have a role to play in cancers like its neighboring gene THAP9-AS1^11^ and other THAP family proteins like THAP1^65^ & 11^66, 67^. Moreover, it is also reported that hTHAP9 shows 5-fold upregulation in TBM (Tuberculous Meningitis) patients co-infected with HIV compared to patients with TBM^68^.

**Figure 15:**
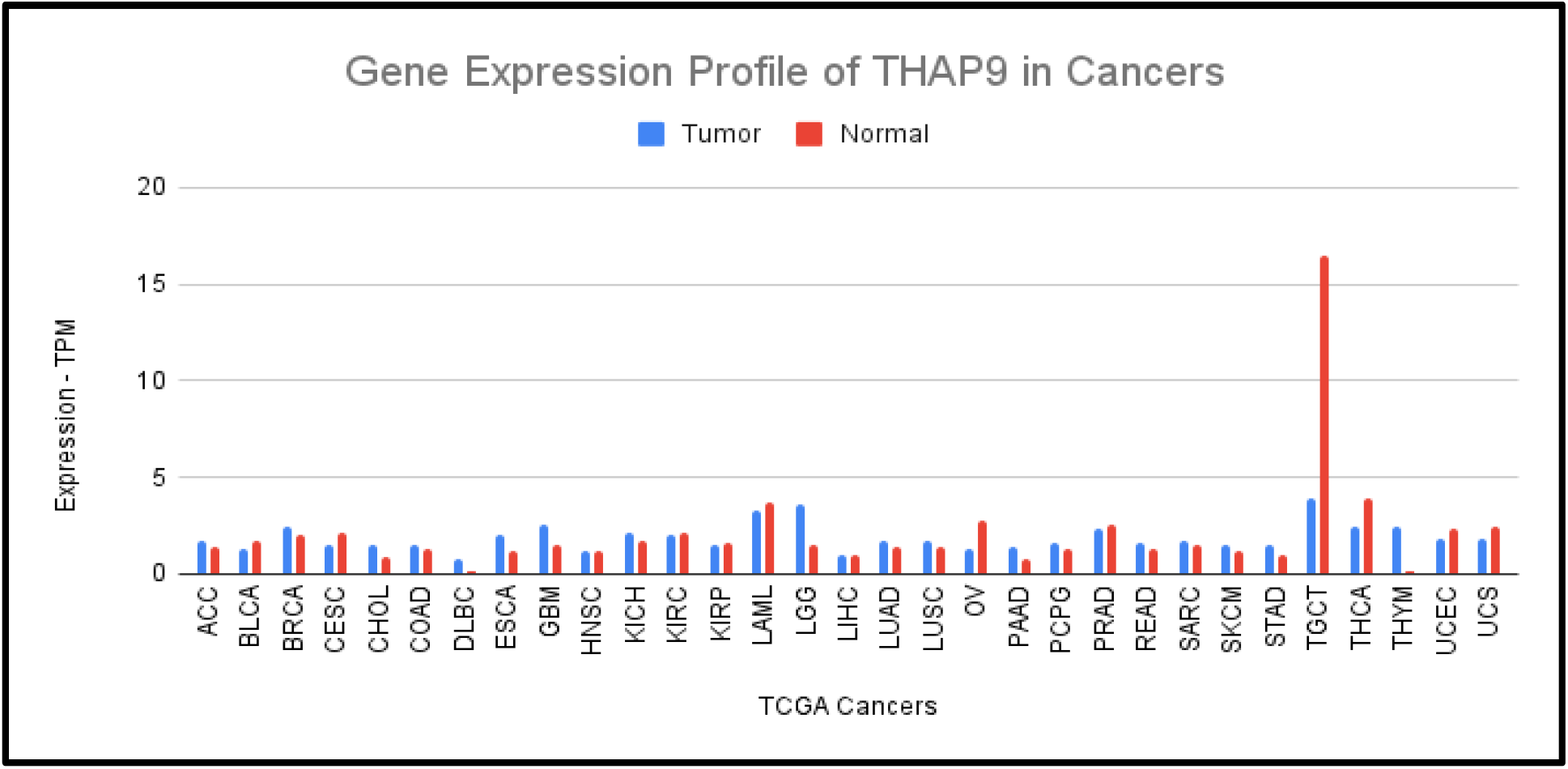
Gene Expression Profile of THAP9 in Cancers. The pan-cancer expression profile of THAP9 has been plotted using the GEPIA tool, which used cancer and matched normal from TCGA and GTEx portal. Pan-cancer view of THAP9 expression showed overexpression of THAP9 in Testicular Cancer (TGCT) and reduced expression of THAP9 in Thymic Epithelial Tumors (THYM). ACC, adrenocortical carcinoma; BLCA, bladder urothelial carcinoma; BRCA, breast invasive carcinoma; CESC, cervical and endocervical cancers; CHOL, cholangiocarcinoma; COAD, colon adenocarcinoma; DLBC, lymphoid neoplasm diffuse large B-cell lymphoma; ESCA, esophageal carcinoma; GBM, glioblastoma multiforme; HNSC, head and neck squamous cell carcinoma; KICH, kidney chromophobe; KIRC, kidney renal clear cell carcinoma; KIRP, kidney renal papillary cell carcinoma; LAML, acute myeloid leukemia; LGG, brain lower grade glioma; LIHC, liver hepatocellular carcinoma; LUAD, lung adenocarcinoma; LUSC, lung squamous cell carcinoma; MESO, mesothelioma; OV, ovarian serous cystadenocarcinoma; PAAD, pancreatic adenocarcinoma; PCPG, pheochromocytoma and paraganglioma; PRAD, prostate adenocarcinoma; READ, rectum adenocarcinoma; SARC, sarcoma; SKCM, skin cutaneous melanoma; STAD, stomach adenocarcinoma; STES, stomach and esophageal carcinoma; TGCT, testicular germ cell tumors; THCA, thyroid carcinoma; THYM, thymoma; UCEC, uterine corpus endometrial carcinoma; UCS, uterine carcinosarcoma; UVM, uveal melanoma.

## Discussion

Human THAP9 (hTHAP9) is a transposable element-derived gene that encodes the hTHAP9 protein, which is a homolog of Drosophila P-element transposase (DmTNP). THAP9 possesses a C2CH type DNA binding THAP domain which is shared between THAP domain-containing proteins (human THAP proteins, CDC14, CTBP1, Lin36, Lin15B, etc.), DmTNP, and zebrafish Pdre2^3, 69, 70^. P element-like transposable elements or P element transposase-like genes are present in other eukaryotes including the sea squirt Ciona, sea urchin, and hydra^71, 72^.

We observed that the orthologs of THAP9 (in 216 species) revealed remarkable conservation across species, although only 1 positively selected site was identified. Across all ortholog species we analyzed in this study, THAP9 has been subjected to a notable overall negative selection, with branch ω-values like those typically observed for housekeeping and essential genes^73^. Further supporting this notion, hTHAP9 is placed in head-2-head architecture along with the THAP9-AS1 gene^74^, which is a commonly reported architecture amongst the housekeeping genes in mammals^75^. According to NCBI, in addition to the studied orthologs, THAP9 also has homologs in chimpanzees, Rhesus monkeys, dogs, cows, chickens, and frogs. In the future, it will be interesting to compare the THAP9 orthologs and homologs and also investigate the role of THAP9 in organisms that have both THAP9 homolog and ortholog sequences.

The negative selection, likely a response to strict functional constraints on THAP9, has yielded a high degree of conservation, not only at the sequence level but also at the domain organization level. Generally, proteins are more conserved at the tertiary structure level than at the amino acid sequence level^7^ and random mutations and selection restrictions help in the structural and functional conservation of protein domains^77, 78^. Interestingly, when comparing the THAP9 orthologs, it was observed that the zinc-coordinating C2CH motif and base-specific DNA-binding residues of the THAP domain were highly conserved. On the other hand, class-specific distinct patterns were observed in other domains. For example, certain functional residues like the catalytic D374(in hTHAP9) in the RNaseH-like domain, aligned with Glycine in all Avian orthologs. The Leucine Zipper region also had 4 more conserved Leucine other than 4 Leucine forming the L-Zipper Region. Moreover, the Leucine-Zipper, bipartite NLS, and other functional motifs present in all mammalian orthologs were absent in other classes, thus implying their possible functional importance in mammals.

We identified several disordered binding regions (DBR) in hTHAP9 protein (**Supplementary Table 1**) that are typically involved in protein-protein interactions ^79^. The DBR region with the highest IUPRED^80^ score is between residues 1-180 and overlaps with the THAP domain and L-Zipper region. The DBRs could also facilitate intramolecular interactions within THAP9 via motifs^81^. Moreover, predicted short linear motifs (**Supplementary Table 2**) in hTHAP9 may be involved in binding regulatory molecules like casein kinase 1 and 2 orHCF1.

The physiological role of hTHAP9 and its disease associations still remain largely unexplored. THAP9-AS1^11^ (the 5’ neighboring gene of THAP9) and other THAP family proteins like THAP1^65^ & 11^66, 67^ have been implicated in various cancers. Thus, we analyzed the SNPs and cancer-associated mutations of hTHAP9 as well as its expression pattern in various cancers and observed its reduced expression in testicular cancer (**Figure. 15**). It is interesting to note that transposition of P-element, the well-studied homolog of hTHAP9, causes hybrid dysgenesis leading to male sterility in Drosophila^82^.

---

**Supplementary Figure 1:**
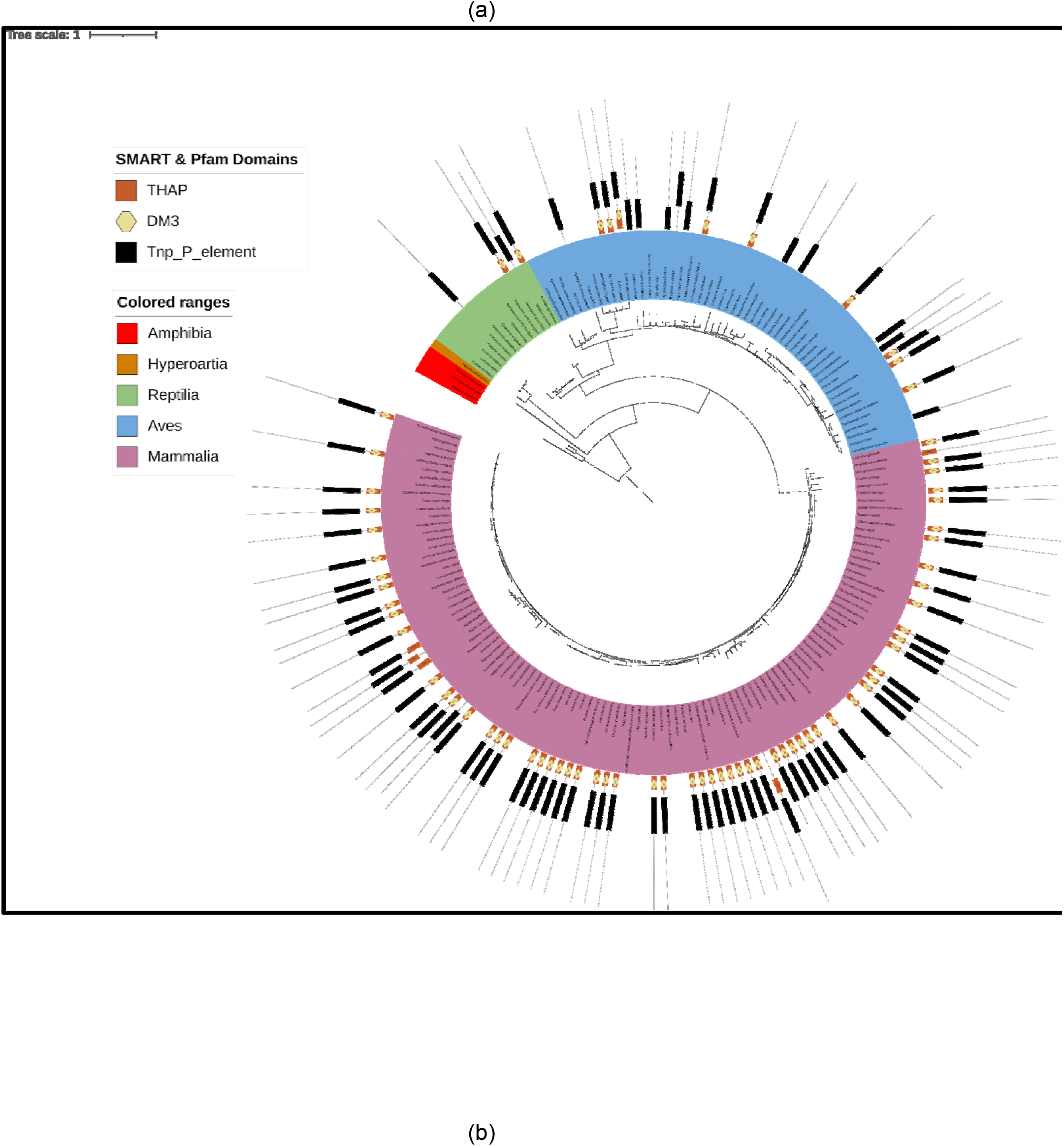
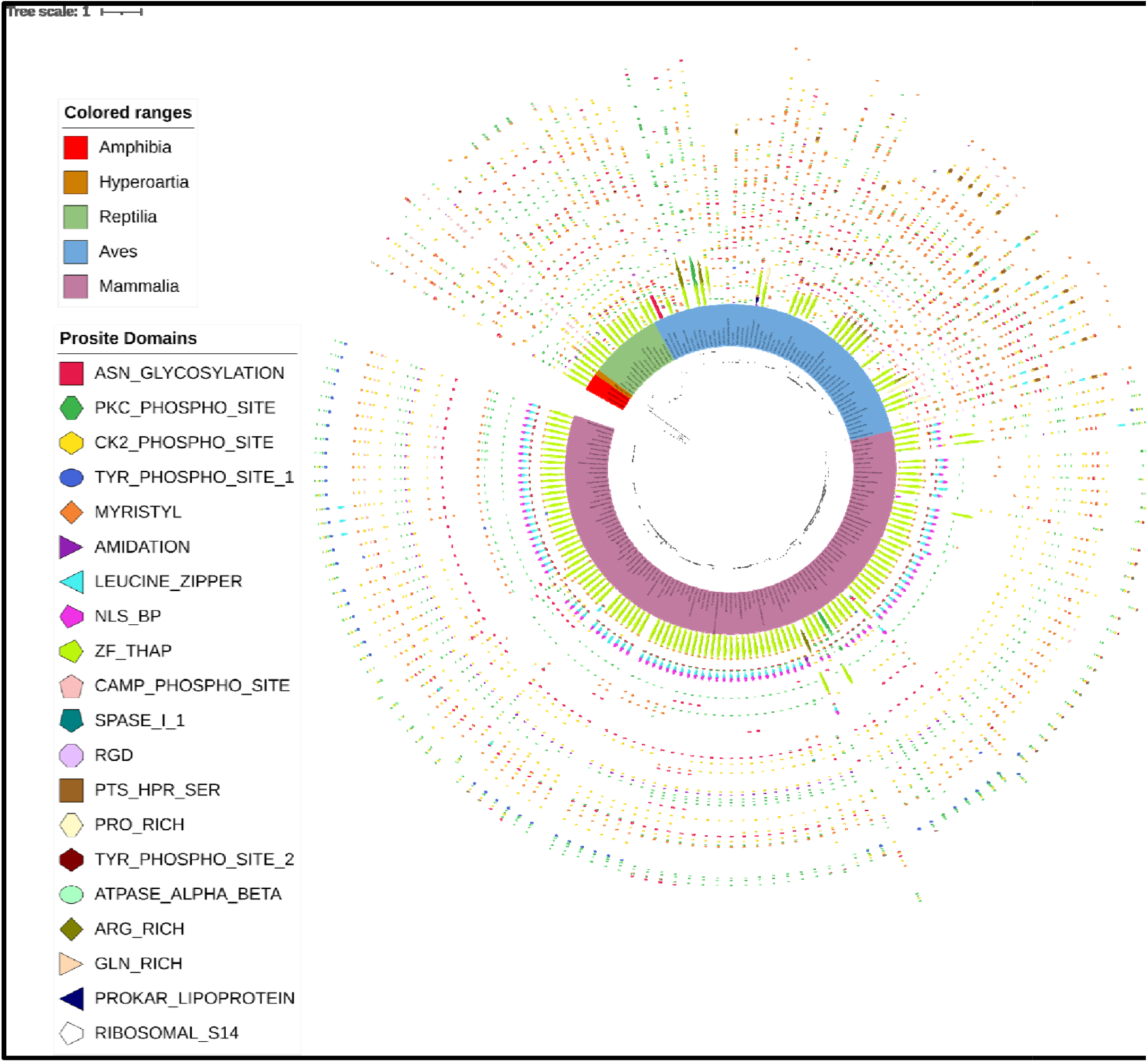
**(a) Full tree representation of previously annotated SMART & Pfam domains of THAP9 orthologs.** Results were visualized on a phylogenetic tree using iTOL. Domain architectures were displayed using the data export feature of SMART. **(b) Full tree representation of newly predicted features of THAP9 orthologs** using Eukaryotic Linear Motif (ELM) and ScanProsite.

## Notes

https://docs.google.com/document/d/1ZAyGVEJ-gKrs_BVDXlEWmEYv8vrRjox_VKzx-zXYEJE/edit?usp=sharing

https://docs.google.com/document/d/1OzIeTtvWl0sGau-g40C0CjQMAwe91VtZY0-UzwA5wRs/edit?usp=sharing

https://drive.google.com/drive/folders/1GF0-aLEpVPhqu1Ioms6XyRq9te-8hpzL?usp=sharing

https://drive.google.com/drive/folders/1mdL71YY4vfffMoVyhOeI0cWXplj339jB?usp=sharing

### Competing Interest Statement

The authors have declared no competing interest.

