## Supplementary figures and images for "Evolutionary analysis of THAP9 transposase: conserved regions, novel motifs"

### Supplementary Figure 1(a)

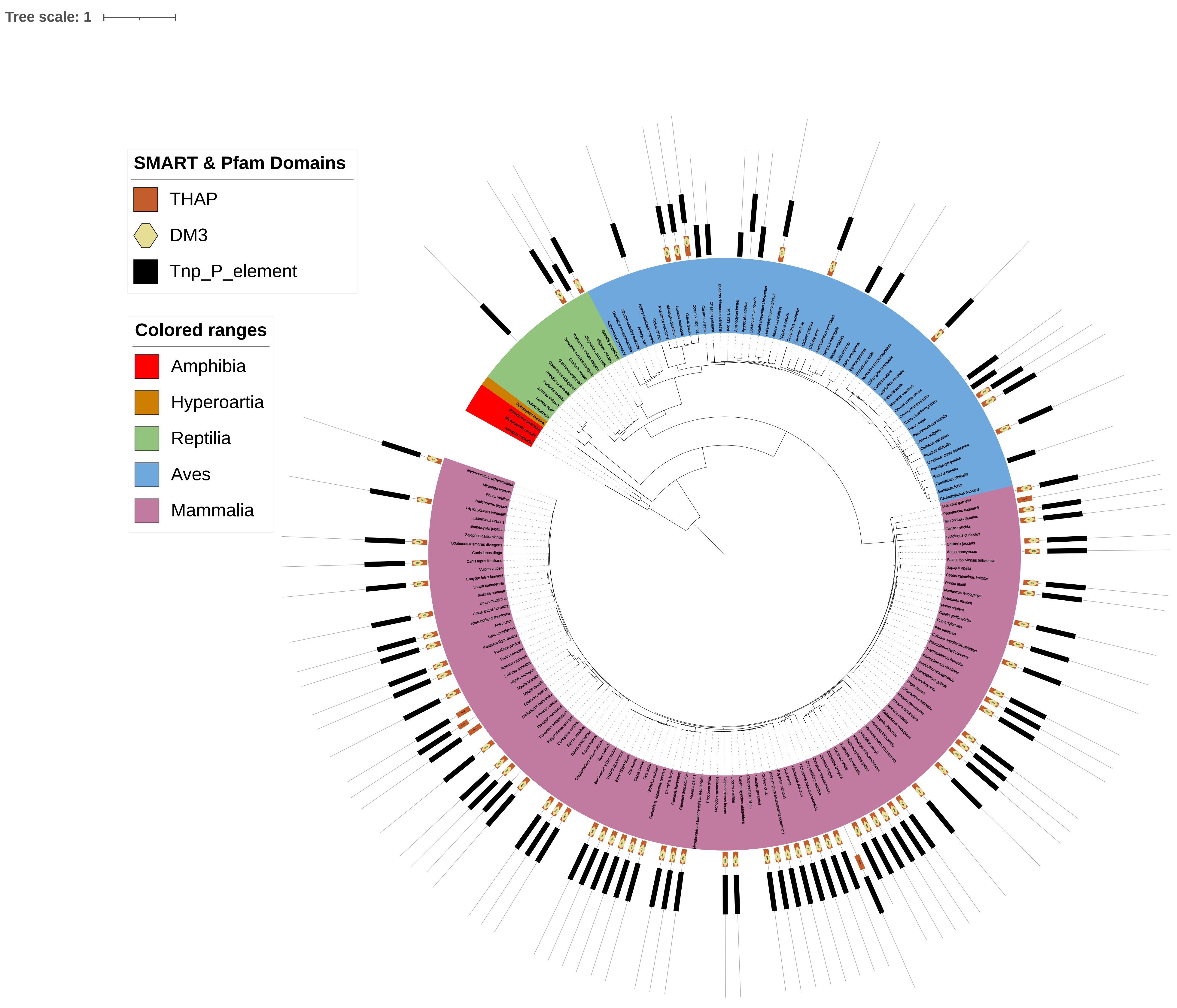

### Supplementary Figure 1(b)

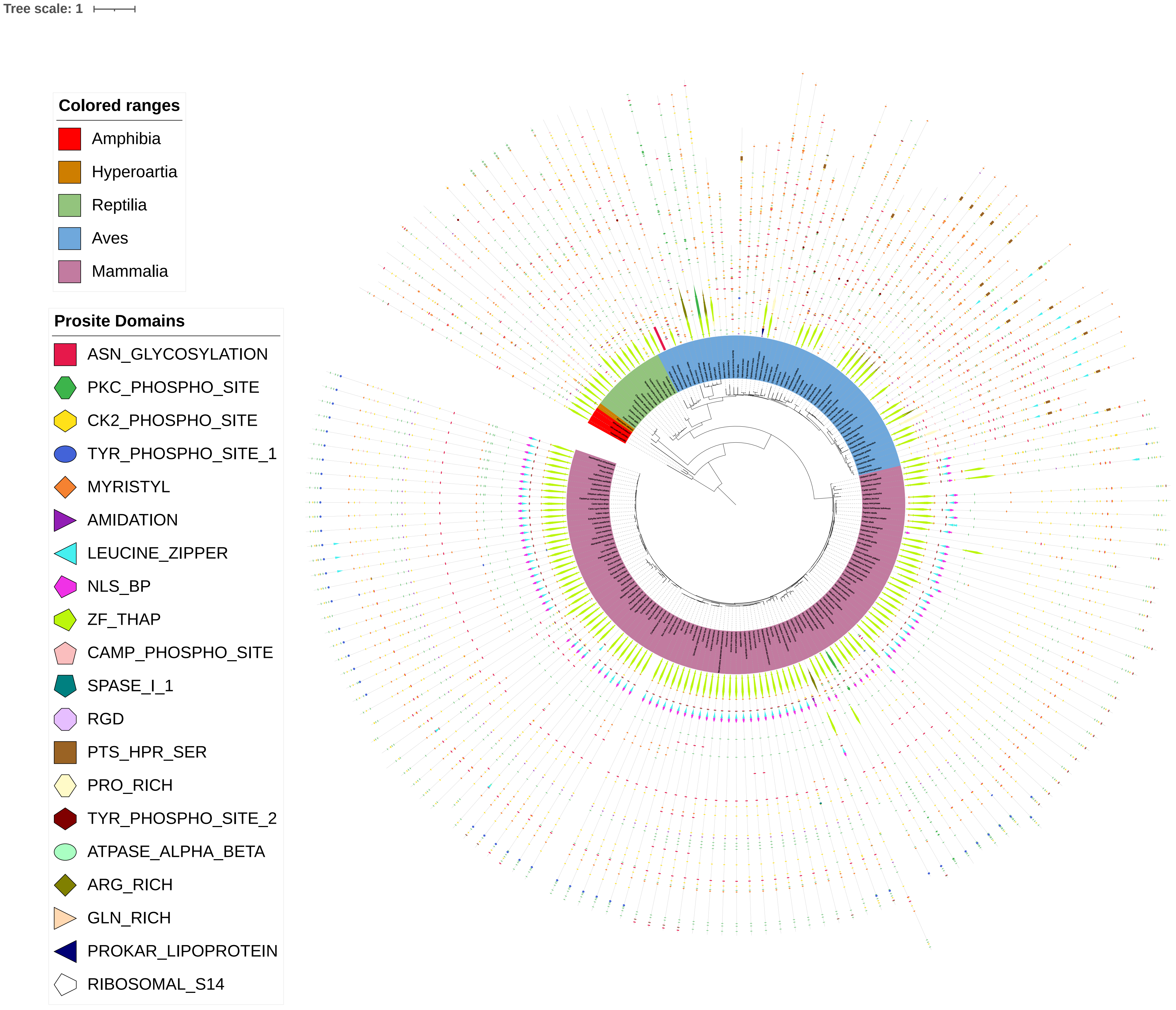
